# Granuloma Dual RNA-Seq Reveals Composite Transcriptional Programs Driven by Neutrophils and Necrosis within Tuberculous Granulomas

**DOI:** 10.1101/2025.04.26.650783

**Authors:** Gopinath Viswanathan, Erika J. Hughes, Mingyu Gan, Ana María Xet-Mull, Graham Alexander, Devjanee Swain-Lenz, Qingyun Liu, David M. Tobin

## Abstract

Mycobacterial granulomas lie at the center of tuberculosis (TB) pathogenesis and represent a unique niche where infecting bacteria survive in nutrient-restricted conditions and in the face of a host immune response. The granuloma’s necrotic core, where bacteria reside extracellularly in humans, is difficult to assess in many experimentally tractable models. Here, using necrotic mycobacterial granulomas in adult zebrafish, we develop dual RNA-seq across different host genotypes to identify the transcriptional alterations that enable bacteria to survive within this key microenvironment. Through pharmacological and genetic interventions, we find that neutrophils within mature, necrotic granulomas promote bacterial growth, in part through upregulation of the bacterial *devR* regulon. We identify conserved suites of bacterial transcriptional programs induced only in the context of this unique necrotic extracellular niche, including bacterial modules related to K^+^ transport and *rpf* genes. Analysis of *Mycobacterium tuberculosis* strains across diverse lineages and human populations suggests that granuloma-specific transcriptional modules are targets for bacterial genetic adaptation in the context of human infection.

**Summary sentence:** Dual host-pathogen transcriptional profiling defines granuloma-specific programs during mycobacterial infection.

## Introduction

Tuberculosis (TB) granulomas represent a distinct environment that facilitates complex interactions between diverse immune cell types and the pathogen *Mycobacterium tuberculosis*, shaping the trajectory of infection. These interactions trigger cellular reprogramming events in the host, including macrophage epithelioid transformation, altered inflammatory responses, and foam cell formation as well as metabolic shifts in both host and pathogen (*1, 2*). The necrotic core of the granuloma is a robust source of host-derived lipids for the bacteria but also exposes the pathogen to challenging conditions for bacterial survival and growth including hypoxia, nutrient limitation, and oxidative stress (*1*). As a highly adapted pathogen, *M. tuberculosis* has evolved mechanisms to survive these conditions (*3*), although the transcriptional reprogramming events driving these adaptations are not fully understood. Studying these host-pathogen dynamics and the underlying transcriptional responses in granulomas is challenging, as most standard animal models do not completely reflect the known features of human TB granulomas, such as central necrosis and epithelioid transformation of macrophages (*4*). In addition, the optical opacity of murine and macaque models of TB infection presents challenges for high-resolution live imaging of host-mycobacterial interactions. Techniques like positron emission tomography-computed tomography and intravital imaging can be used in some of these models to visualize granuloma dynamics, but they present limitations to resolution or are not compatible with continuous long-term imaging (*5–9*).

Neutrophils, the most abundant immune cell types in circulation, had been reported to be recruited to TB granulomas as early as 1931 (*10*), but they remain a relatively understudied cell type in comparison to other immune cells involved in TB pathogenesis (*11*). Although murine models of TB infection initially suggested some host protective functions for neutrophils (*12, 13*), they likely play a predominant host detrimental role as well (*14, 15*). Neutrophils can compromise host resistance to mycobacterial infection through multiple mechanisms including type I interferon-driven NETosis, which leads to tissue destruction, and by creating a nutritionally permissive niche that supports *M. tuberculosis* growth (*14–18*). These dual roles may correspond to discrete time points in infection; in zebrafish larvae, neutrophils can play an initial host protective role during the early stages of mycobacterial infection through oxidative and nitric oxide-dependent killing (*19, 20*). Human studies too present a complex picture of neutrophils in TB. The risk of TB infection was initially found to be inversely associated with peripheral blood neutrophil counts in contacts of patients with active TB (*21*). However, whole blood transcript analysis of active TB patients shows a dominant neutrophil-driven type I interferon signature, which associates with worse outcomes and is predictive of active TB (*22*). Neturophil interactions with other immune cell types and mycobacteria within the mature granulomatous niche have been challenging to observe *in vivo* as have specific bacterial transcriptional alterations that are neutrophil dependent.

Here, we utilized the adult zebrafish - *M. marinum* infection model and a granuloma explant model to study neutrophil dynamics and functions during a critical phase of infection: that of mature necrotic granulomas and paired these analyses to dual host-pathogen transcriptional profiling. Pharmacological or cell-specific genetic inhibition of neutrophil function resulted in a reduction in mycobacterial burden, revealing a pathogen-permissive role within necrotic granulomas. High-resolution microscopy via the Mycobacterial Granuloma Explant Model (Myco-GEM) enabled the visualization of complex interactions between neutrophils, macrophages, and mycobacteria in specific granuloma compartments, suggesting key points in an infectious cycle at which neutrophils mediate disease outcome.

Dual RNA-seq in granuloma samples presents technical challenges, as bacterial transcripts are significantly underrepresented compared to host transcripts in total RNA isolated from these granulomas. To address this issue, we developed an enrichment protocol that leverages compositional differences between host and bacterial mRNA, allowing for the enrichment of bacterial transcripts. Using this granuloma dual RNA-seq strategy, we uncovered a neutrophil-induced mycobacterial transcriptional module that promotes pathogen growth within necrotic granulomas, as well as roles for neutrophils in modulating host inflammation and oxidative stress responses.

Combination of high-resolution imaging and dual transcriptional profiling during this unique stage of infection revealed a previously undefined necrotic granuloma-specific repertoire of genes and networks deployed by pathogenic mycobacteria to support chronic infection. These analyses provide additional insights into the metabolic and physiological states of mycobacteria during their persistence within the granuloma, both in the presence and absence of neutrophils. We identified genes uniquely induced in necrotic granulomas with specific roles in ion scavenging, oxidative stress response, nitrogen metabolism, and lipid metabolism, revealing key mycobacterial transcriptional programs that enable the pathogen to endure and thrive under the harsh conditions present in these structures. Finally, genomic and evolutionary analysis of *M. tuberculosis* lineage-defining variants in diverse clinical strains revealed prominent signatures of evolutionary selection at these bacterial genes in human populations.

## Results

### Granuloma microenvironment associates with neutrophil form and function

TB granulomas display remarkable diversity and can be classified based on their characteristics into cellular/early non-necrotizing, necrotizing caseated, cavitating, or calcified granulomas (*23*). They can also exhibit intrinsic heterogeneity where organized microenvironments can be found within individual granulomas harboring immune cell types with distinct inflammatory statuses, suggesting a role for these compartments in influencing immune cell behaviors (*24*). We aimed to study neutrophil dynamics and their interactions with mycobacteria and other immune cells in a physiologically relevant setting that mimics the complex heterogeneity and spectrum observed in human granulomas. Upon infection with *M. marinum,* adult zebrafish form cellular and necrotizing caseated granulomas resembling the progressive granuloma subtypes found in humans (*25*). We previously established an ex vivo granuloma culture technique called Myco-GEM, wherein organized granulomas from infected adult zebrafish can be microdissected and maintained in a three-dimensional culture (*4*). Here, we utilized this explant system to study the dynamics of fluorescently labeled neutrophils in granulomas isolated from Tg(*lyz:egfp*)*^nz117^* animals (*26*).

First, we microscopically assessed these granuloma explants to ensure their heterogeneity and indeed observed diversity among them. Based on the bacillary distribution pattern and their physical characteristics, we classified them into two types (Fig. 1A). In type I granulomas, *M. marinum* is mostly contained within the necrotic core, which is surrounded by well-formed epithelioid macrophages with very few or no extra-necrotic bacilli, resembling caseating necrotic lesions. In type II granulomas, *M. marinum* can be seen in the extra necrotic region (ENR) in addition to the necrotic core. Compared to the abundant mycobacterial growth in the necrotic core, the bacterial burden in the ENR was lower, with individual bacilli more frequently observed in this region. In the bright-field images, ENR looked different than the necrotic core and appeared to have cellular layers. Hence, the type II granulomas are likely hybrid structures in which *M. marinum* can be found both in necrotic and cellular regions (Fig. 1A, B).

**Fig. 1.**
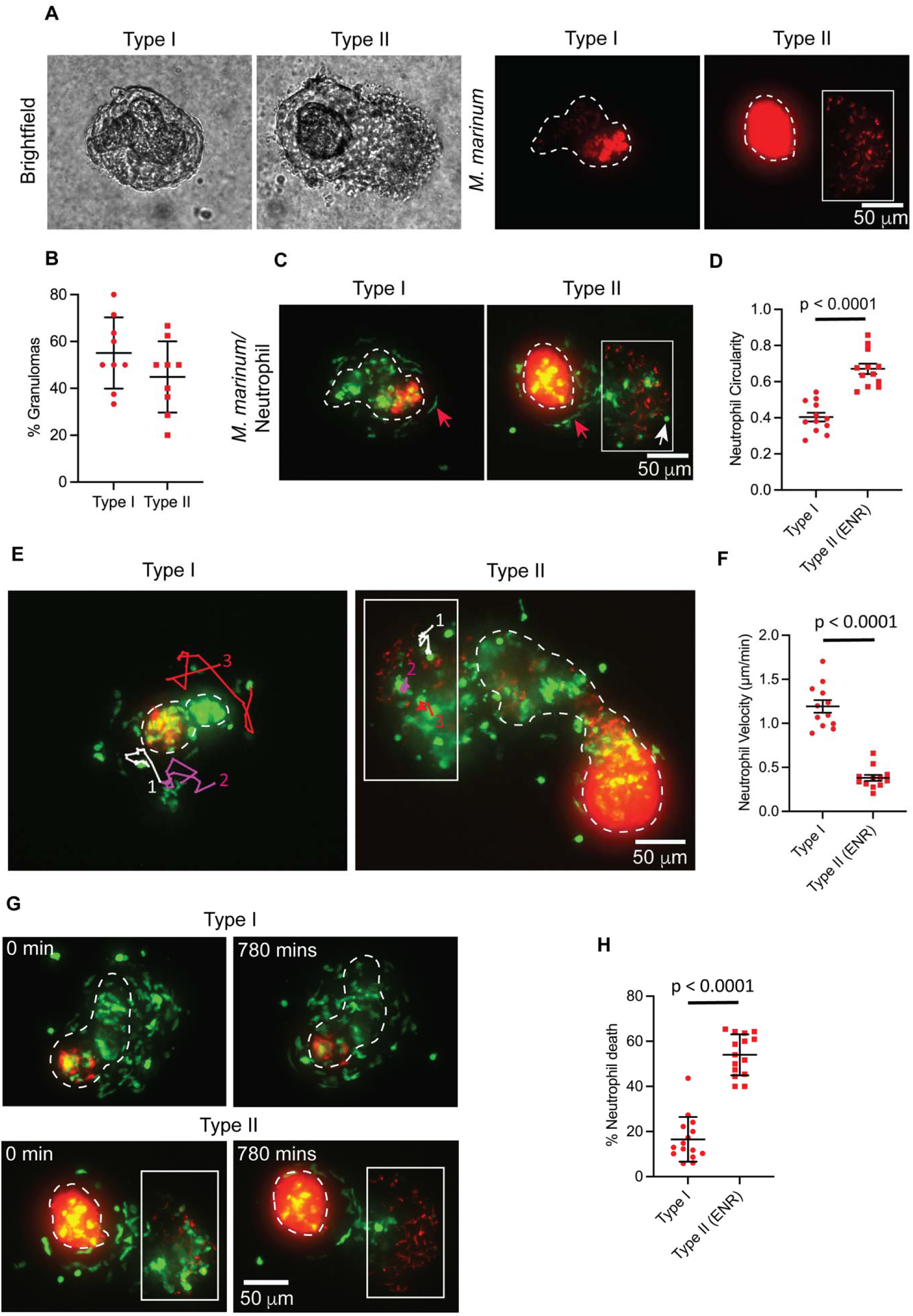
Neutrophil morphology, dynamics, and survival kinetics vary by granuloma subtype. **A)** Representative bright-field and fluorescent images of granuloma explants from adult zebrafish infected with *M. marinum* (red). Granulomas were categorized into two subtypes based on their bacillary distribution pattern. White dashed lines mark necrotic core boundaries. *M. marinum* present outside the necrotic core in type II granulomas (Extra Necrotic Region (ENR)) are indicated by a white box. **(B)** Mean percentage of granuloma subtypes observed in granuloma explants. Error bars show standard deviation (SD). Each data point represents the percentage of granuloma subtype from a single experiment. n = 9 independent experiments, 91 total granuloma explants from 27 wild-type animals were analyzed. **C)** Fluorescent images of granuloma explants from **(A)** showing altered neutrophil morphologies in granuloma subtypes. Neutrophils are shown in green. Red arrows indicate elongated neutrophils; white arrows indicate rounded neutrophils. **(D)** Neutrophil shapes in type I and ENR of type II granulomas, represented as mean circularity with error bars indicating standard error of the mean (SEM). Each data point indicates mean neutrophil circularity from a single granuloma, with a maximum of 1 indicating completely circular neutrophils. 306 neutrophils from 12 type I granulomas and 183 neutrophils from the ENR of 12 type II granulomas were analyzed. Statistical comparison by two-tailed, unpaired t-test. **E)** Neutrophils tracked for a period of 230 mins in type I and ENR of type II granulomas. Individual tracks are labeled white, magenta, and red and numbered. **(F)** Mean neutrophil velocities in granuloma subtypes with error bars indicating SEM. Each data point denotes the mean velocity of 5 neutrophils tracked for 230 mins from a single granuloma. 60 neutrophils from 12 granulomas were tracked for each granuloma subtype. Statistical comparisons by unpaired t-test with Welch’s correction. **(G)** Fluorescent time-lapse images showing neutrophil viability in type I and ENR of type II granulomas observed over 780 mins. **(H)** Mean percentage of neutrophil death observed in granuloma subtypes with error bars denoting SD. Each data point represents the percentage of neutrophil death observed for 780 mins in a single granuloma, n = 15 granulomas from each subtype; type I – 1080 neutrophils analyzed; type II (ENR) - 532 neutrophils analyzed. Two-tailed, unpaired t-test was used. **(D-H)** Data pooled from three independent experiments with a total of 9 animals. Fluorescent images are 100 μm maximum projections. Scale bar, 50 µm.

We then investigated whether the diverse microenvironment within these heterogenous granulomas influenced neutrophil status and found that the neutrophils exhibited varied morphologies across granuloma types. In type I granulomas, neutrophils were primarily elongated, whereas in type II granulomas, they exhibited a dimorphic pattern, with elongated neutrophils appearing near the necrotic core and rounded neutrophils in the *M. marinum* containing ENR (Fig. 1C). Neutrophils in the ENR of type II granulomas were significantly more circular than those in type I granulomas (Fig. 1D). Neutrophils, in general, are known for their remarkable deformability, and alterations in their morphology are often associated with behavioral changes (*27–30*). We performed high-resolution long-term live imaging of explant granulomas tracking fluorescent neutrophils and found that elongated neutrophils in type I granulomas were highly dynamic. They had reversible access to different regions in the granulomas and were found to be patrolling the entire cellular layer surrounding the necrotic core (Fig. 1E and Movie S1). Elongated neutrophils near the necrotic core of type II granulomas behaved similarly to that of their type I counterparts (Movie S1). In contrast, rounded neutrophils in the ENR of type II granulomas were found to be less motile and remained confined to this region (Fig. 1E and Movie S1). These observations were further validated by velocity measurements (Fig. 1F). The dimorphic neutrophils in type II granulomas were largely segregated within their respective microenvironments. However, we regularly observed migration of elongated neutrophils present near the necrotic core towards *M. marinum* containing ENR (Fig. S1 and Movie S2). From these observations, we propose that the dynamic, elongated neutrophils may perform a patrolling function and are capable of intercompartmental movements in type II granulomas in response to possible chemotactic stimuli. These neutrophil subtypes also displayed altered viability, with elongated neutrophils exhibiting a prolonged lifespan in type I granulomas, while rounded neutrophils in the type II ENR showed rapid cell lysis (Fig. 1, G, H and Movie S3). Neutrophils are known to undergo necrosis or programmed cell death upon infection with *M. tuberculosis* (*31, 32*). Hence, we looked at the infection status of these rapidly lysing neutrophils in the type II ENR. We noticed both infected and uninfected neutrophils undergoing cell death in this region, suggesting that, in addition to infection status, the microenvironment in type II ENR may affect their viability (Fig. S2 and Movie S4).

Given that macrophages are the major cell type within granulomas (*2*), we next decided to investigate their interactions with neutrophils using a double transgenic line (Tg(*lyz:egfp*)*^nz117^*; Tg(*irg1:tdTomato*) *^xt40^*), where neutrophils and macrophages are differentially labeled using the *lyz* and *irg1* (also called *acod1*) promoters respectively (*26, 33*). We found that the macrophages are the predominant infected cell type in the ENR of type II granulomas, and like neutrophils, they appeared to undergo cell lysis events, releasing bacilli into the extracellular milieu (Fig. S3A and Movie S5). The lysis of both neutrophils and macrophages in the ENR suggests that these could be transitional regions evolving into necrotic lesions, potentially leading to the formation of multicentric granulomas with multiple necrotic cores, as reported previously (*25, 34*). In contrast, macrophages in type I granulomas displayed a longer lifespan (Fig. S3A and Movie S5). We observed transient interactions between the dying neutrophils and the infected macrophages in the ENR (Fig. S3B and Movie S5). Nearby macrophages made unsuccessful attempts to phagocytose the extracellular bacteria released from the infected macrophages, indicating that these bacteria are largely inaccessible to the nearby phagocytes for reinfection (Fig. S3B and Movie S5). To further analyze these interactions, we examined tissue sections from *M. marinum-*infected double-transgenic animals at higher magnification. In addition to the cellular contacts between neutrophils and infected macrophages, we observed both infected and uninfected macrophages scavenging dying neutrophils and smaller cellular debris from lysed neutrophils within the type II ENR (Fig. S3C). This phenomenon of neutrophil efferocytosis by macrophages has been previously reported to control mycobacterial growth in *in vitro* coculture settings (*31, 35*), but its role in the context of granulomas remains unclear. Furthermore, the heterogeneity in neutrophil morphologies observed in granuloma explants (Fig. 1C) was also evident in granuloma-containing tissue sections (Fig. S3C), confirming that these observations were not an artifact of Myco-GEM culture conditions. Together, these results suggest that specific regions within granulomas serve as hotspots for complex, multi-faceted neutrophil interactions with mycobacteria and macrophages.

### Single-cell RNA-seq analysis identifies transcriptional heterogeneity in granuloma neutrophils

Previously, we performed single-cell RNA-seq (scRNA-seq) analysis of zebrafish granulomas to investigate their cellular composition and diverse inflammatory status. This led to the identification of distinct cell clusters in granulomas, each exhibiting transcriptional signatures corresponding to different cell types, including neutrophils (*36*). To determine whether the functional heterogeneity of granuloma neutrophils is reflected in their transcriptional profiles, we performed a sub-clustering analysis of the scRNA-seq data from 256 granuloma neutrophils (Table S1) from our prior dataset (*36*). Based on their unique expression profiles, granuloma neutrophils were classified into three subgroups, with a majority of them falling under group 0, followed by equal representation of group 1 and group 2 neutrophils (Fig. S4A and Table S1). Gene ontology analysis of the differentially expressed genes (with a p_adj_ cutoff value of less than 0.1) in group 0 neutrophils revealed an enrichment of transcripts (*b2m, mhc2dab, cd74b*) encoding major histocompatibility complex (MHC) class I and class II components within this subset (Fig. S4B, E (i), Table S1 and Table S2). Although neutrophils are not primarily recognized as conventional antigen-presenting cells, chronic inflammatory conditions, such as those observed in rheumatoid arthritis, are known to induce MHC-II expression in neutrophils, promoting their antigen-presenting capacity and interactions with T cells (*37*). Notably, like these group 0 neutrophils, a neutrophil subset identified in Cynomolgus macaque TB granulomas was found to specifically express MHC-II components, suggesting that this subpopulation may be conserved across species in the context of chronic TB infection (*38, 39*).

Group 1 neutrophils were enriched with transcripts associated with key processes such as actin filament binding and organization (*capgb, flna*, *pfn1, cotl1*), Arp2/3 actin nucleation complex (*arpc2*, *arpc3*), neutrophil migration (*s1pr4*, *rac2*) and glycolysis (*gapdh*, *gpia, pgk1*) (Fig. S4C, E (ii), Table S1 and Table S2). This suggested that this neutrophil subtype might be actively migrating cell population with their metabolism tuned towards glycolysis to support their function. Their transcript profile aligns with the microscopic observation of a highly dynamic, elongated neutrophil subpopulation actively patrolling the granulomas (Fig. 1E and Movie S1), indicating that these transcripts may drive distinct morphology and motile behavior. Further, Group 1 neutrophils shared genes related to actin filament binding and organization, as well as the Arp2/3 nucleation complex, such as *pfn1, cotl1, capgb,* and *arpc2,* with a neutrophil subcluster from Cynomolgus macaque granulomas, suggesting potentially conserved functions (*38*).

Group 2 neutrophils showed enrichment of transcripts associated with granulocyte differentiation, such as *cebpa* and *csf3a*, as well as the pro-inflammatory chemokine *cxcl11.1* (*36*), reflecting that this subgroup may be relatively more mature and primed for effector functions (Fig. S4D, Table S1 and Table S2). This subset also expressed pro-apoptotic genes including *nr4a1*, *pmaip1* (NOXA), *anxa11a*, and *pdcd6* (ALG2) (Table S1, Table S2 and Fig. S4E (iii)), which act by modulating the functions of apoptotic regulators such as BCL-2 and MCL-1 (*40–43*). These findings correlate with our observation of the dying neutrophil population in the ENR of type II granulomas (Fig. 1G and Movie S3). The combined expression of cytokine, chemokine, and apoptotic markers in this neutrophil population implies that they may participate in a pro-inflammatory response while being highly responsive to apoptotic stimuli. Collectively, these data suggest a transcriptional basis for the microscopically observed functional heterogeneity in granuloma neutrophils and point to the potential conservation of functions in neutrophil subgroups across non-human primate granulomas.

### Neutrophils play a pathogen-permissive role in necrotic granulomas

Our ability to culture and image intact necrotic granulomas *ex vivo* using Myco-GEM allowed us to chemically target pan-neutrophil populations within the granulomas and assess their acute impact on mycobacterial burden. We treated the granuloma explants with Duvelisib, a U.S. FDA-approved orphan drug used for the treatment of refractory chronic lymphocytic leukemia and small lymphocytic lymphoma (*44*). Duvelisib targets both the phosphoinositide 3-kinase (PI3K) isoforms γ and δ, which are crucial regulators of key neutrophil functions, including actin cytoskeleton remodeling, chemotaxis, superoxide generation, and degranulation (*44, 45*). Consistent with its reported activities, Duvelisib treatment altered neutrophil morphology and dynamics within granulomas, resulting in increased circularity and reduced motility compared to DMSO-treated controls (Fig. 2, A, B, C, D and Movie S7). These changes suggested impaired neutrophil functionality in Duvelisib-treated granulomas. We then evaluated the effect of neutrophil inhibition on granuloma bacterial burden by measuring the mycobacterial fluorescence four days post-treatment with Duvelisib. This method has been previously used to reliably quantify bacterial burden in granuloma explants (*46*). Using this approach, we observed a significant reduction in mycobacterial burden in Duvelisib-treated granulomas compared to the DMSO-treated control, indicating that neutrophils support mycobacterial growth in necrotic granulomas (Fig. 2, E and F). Importantly, Duvelisib treatment did not affect the growth of *M. marinum* in broth culture (Fig. 2G), suggesting that the reduction in bacterial burden is mediated through its impact on neutrophils rather than a direct bactericidal effect.

**Fig. 2.**
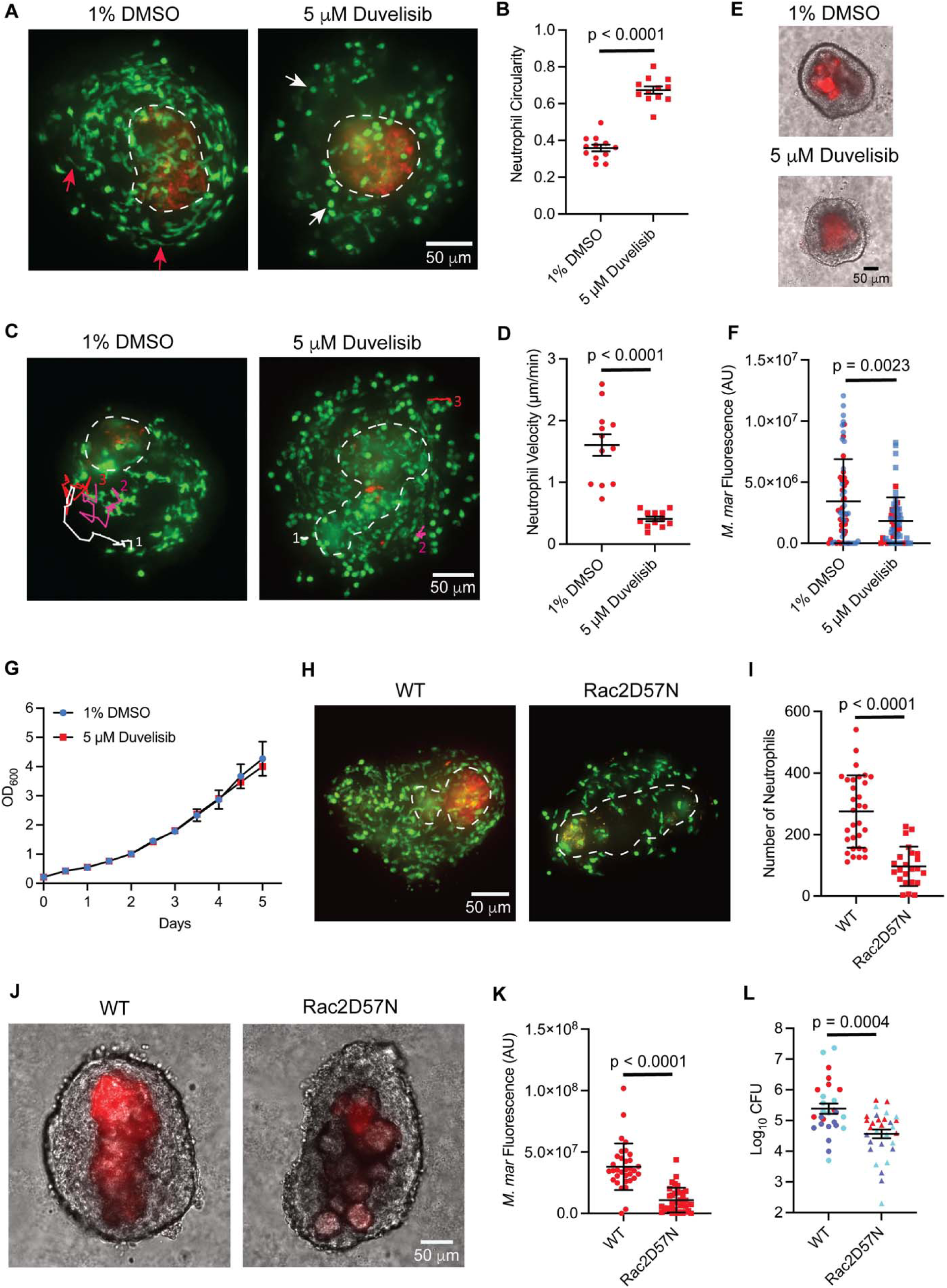
Pharmacological and genetic interventions of granuloma neutrophil functions lead to the reduction in mycobacterial burden. **(A)** Fluorescent images showing altered neutrophil morphology in granuloma explants 5 hours post-treatment with duvelisib versus vehicle control. Red arrows - elongated neutrophils in a vehicle-treated granuloma, white arrows – rounded neutrophils in duvelisib-treated granuloma. **(B)** Mean circularity of neutrophils from vehicle-treated and duvelisib-treated granulomas with error bars indicating SEM. Each data point represents the mean circularity of neutrophils from a single granuloma measured 5 hours post-treatment with vehicle/duvelisib. Vehicle – 300 neutrophils from 12 granulomas analyzed, Duvelisib – 389 neutrophils from 12 granulomas analyzed. **(C)** Neutrophils, tracked over 230 mins in granuloma explants 5 hours post-treatment with vehicle/duvelisib. Individual tracks are labeled white, magenta, and red and numbered. **(D)** Mean neutrophil velocities from vehicle-treated and duvelisib-treated granulomas with error bars indicating SEM. Each data point denotes the mean velocity of 5 neutrophils tracked for 230 mins from a single granuloma 5 hours post-treatment with vehicle/duvelisib. 60 neutrophils from 12 granulomas were analyzed for each treatment. **(E)** Representative images of *M. marinum* (red) infected granuloma explants 4 days post-treatment with vehicle/duvelisib. Bright-field and fluorescent channels are merged. (F) *M. marinum* burden in explant granulomas 4 days post-treatment with vehicle/duvelisib represented as mean arbitrary fluorescence units (AU). Error bar indicates SD. Data pooled from two independent experiments, differentiated by red and blue data points. Each data point represents *M. marinum* fluorescence measured in a single granuloma, n = 60 granulomas for vehicle & 62 granulomas for duvelisib treatment, obtained from 6 animals each. No significant differences in bacterial burden were observed before treatment. (G) *in vitro* growth profile of *M. marinum* in the presence of duvelisib/vehicle. Error bars represent SD. **(H)** Fluorescent images showing the number of neutrophils recruited to the Rac2D57N granuloma and its WT sibling. **(I)** Mean number of neutrophils observed in the Rac2D57N granulomas and their WT siblings with the error bars denoting SD. Each data point represents the number of neutrophils observed in a single granuloma, n = 31 granulomas for the WT & 23 granulomas for the Rac2D57N, obtained from 3 animals each; **(J)** Representative images showing *M. marinum* burden in the WT and Rac2D57N granulomas. (K) *M. marinum* burden in WT and Rac2D57N granulomas represented as mean arbitrary fluorescence units (AU). Granulomas were dissected from 14 dpi animals. Error bar indicates SD. Each data point represents *M. marinum* fluorescence measured in a single granuloma, n = 33 granulomas for the WT & 35 granulomas for the Rac2D57N, obtained from 3 animals each; (L) *M. marinum* burden in 14 dpi WT and Rac2D57N fish represented as mean Log_10_CFUs. Error bars show SEM. Data pooled from three independent experiments, differentiated by red, purple, and cyan-colored data points. 27 WT and 30 Rac2D57N fish were used. **(A, C, H – K)** Representative of three independent experiments. **(E, G)** Representative of two independent experiments. **(B, D)** Data pooled from three independent experiments, n = 9 animals in total for each group. **(B, L)** Two-tailed, unpaired t-test, **(D, F, I & K)** Unpaired t-test with Welch’s correction. Fluorescent images are 100 μm maximum projections. Scale bar, 50 µm.

To further validate our findings from Duvelisib treatment and eliminate the possibility of off-target drug effects or effects on other cell types from pharmacological inhibition, we employed a more targeted neutrophil-specific genetic approach. The Rho GTPase Rac2 acts downstream of the PI3K signaling pathway and is positively regulated by this kinase family, with a functionally interdependent role in controlling key neutrophil activities like chemotaxis, ROS production, and degranulation (*47*). Previously, a point mutation in this gene was reported in a patient, substituting Asp57 with Asn in one of the alleles, which caused severe neutrophil dysfunction and increased susceptibility to bacterial infections (*48*). Since this residue and the Rac2 gene are highly conserved in zebrafish, a transgenic animal model was developed to mimic this clinical phenotype in which the dominant-negative version of the cognate gene (Rac2D57N) along with the fluorescent marker mCherry, specifically in neutrophils, driven by the neutrophil-specific promoter *mpx.* The functions of Rac2D57N-expressing neutrophils were found to be severely compromised in this transgenic fish, as they failed to respond to tail wounding and *Pseudomonas aeruginosa* infection (*49*). We crossed this Rac2D57N fish with Tg(*lyz:egfp*)*^nz117^* animals to enable better visualization of neutrophils. This double-transgenic fish was then used to investigate the direct impact of neutrophil inhibition on mycobacterial burden. We first assessed whether the functional impairment of Rac2D57N neutrophils could be reproduced in the context of mycobacterial infection by performing a neutrophil recruitment assay following *M. marinum* infection in the hindbrain ventricle. A significant reduction in the recruitment of Rac2D57N neutrophils to the infection site was observed compared to wild-type neutrophils, indicating their compromised functionality (Fig. S5A and B). We then examined the Rac2D57N neutrophil recruitment to the necrotic granulomas and found a significant reduction in their numbers in comparison to the neutrophils in wild-type granulomas, confirming the recruitment defects during chronic infection (Fig. 2, H, I and Fig. S5C). These poorly recruited Rac2D57N neutrophils in the granulomas often displayed a more circular morphology compared to wild-type neutrophils, similar to what was observed with the Duvelisib treatment (Fig. S5D). Consistent with the findings from Duvelisib treatment, Rac2D57N granulomas exhibited a significantly lower bacterial burden compared to wild-type granulomas, as determined by mycobacterial fluorescence (Fig. 2, J, K and Fig. S5E). This result was further verified through whole-animal CFU measurements in both wild-type and Rac2D57N fish (Fig. 2L). Thus, by combining pharmacological and genetic approaches that target the PI3K-Rac2 signaling axis, we identified a role for neutrophils in supporting mycobacterial growth during chronic infection in necrotic granulomas. These results agree with recent studies in the C3HeB/FeJ mouse model, which develops necrotic lesions and demonstrates a pathogen-permissive role of neutrophils during TB infection (*14, 15, 18*).

### A granuloma dual RNA-seq strategy to decode neutrophil-mycobacterial interactions in necrotic granulomas

The complex three-way interactions between neutrophils, macrophages, and mycobacteria in the granulomas, as observed through live imaging (Fig. 1 and Fig. S3), suggested that neutrophils could drive granuloma-specific phenotypes either by directly influencing bacterial adaptation or modulating host immune responses. To investigate these possibilities, we employed a dual RNA-seq approach to simultaneously capture transcriptional changes in both host cells and *M. marinum* within wild-type and neutrophil-deficient (Rac2D57N) granulomas. Profiling the host transcriptome from granulomas dissected from infected adult zebrafish is a relatively straightforward approach that has been previously used to assess macrophage reprogramming into epithelial-like cells within these structures (*50*). However, the abundance of host transcripts over bacterial transcripts in the total RNA isolated from granulomas presents challenges in achieving adequate coverage of the mycobacterial transcriptome. Recently, bacterial mRNA enrichment protocols have been developed in the context of *M. tuberculosis* mouse infection to identify gene signatures governing in vivo adaptations to alveolar and interstitial macrophages (*51, 52*). These protocols were based on various techniques, including the gentle and specific lysis of the infected host cells in Trizol, which releases *M. tuberculosis,* followed by direct enrichment of the bacteria via centrifugation (*52*). Another method involved the use of a custom biotinylated bait library complementary to the *M. tuberculosis* transcriptome to selectively enrich bacterial transcripts from the sample (*51*). Although highly valuable, these studies were limited to intracellular bacteria, as the C57BL/6J mice used do not develop necrotic granulomas.

Here, we leveraged a fundamental difference between eukaryotic and prokaryotic RNA biology to specifically enrich *M. marinum* transcripts from our samples. Eukaryotic mRNAs are typically polyadenylated at the 3’ end, whereas bacterial RNAs generally lack poly(A) tails. Our approach involved selective depletion of these poly(A)-tail-containing host mRNAs using oligo(dT) beads. This strategy has been previously used to successfully enrich bacterial transcripts from *Chlamydia trachomatis*-infected human HEp-2 epithelial cells (*53*). We first split the total RNA isolated from the dissected granulomas into two parts. One part was subjected to zebrafish and *M. marinum* rRNA depletion using custom-designed species-specific probes, followed by cDNA library preparation and sequencing (Fig. 3A and Table S3). We targeted a sequencing depth of ∼100 million total reads per sample, aiming to adequately capture the *M. marinum* transcriptome. While sufficient reads were obtained for the host transcriptome as expected, the bacterial mRNA reads accounted for only 0.1% - 0.25% of the total reads, which were insufficient for downstream transcriptome analysis (Fig. 3, B and C). Hence, the second portion of the total RNA was subjected to a robust depletion strategy involving treatment with the oligo(dT) beads to specifically deplete poly(A) tail-containing host mRNAs, as discussed above, followed by rRNA depletion (Fig. 3A). Using this approach, we achieved ∼ 4-9 fold enrichment of bacterial mRNA reads compared to the samples that underwent only rRNA depletion (Fig. 3, B and C). This dual RNA-seq strategy enabled us to simultaneously obtain sufficient host and mycobacterial reads to perform transcriptome analysis from necrotic granulomas, allowing us to examine the transcriptional landscapes of wildtype and Rac2D57N granulomas.

**Fig. 3.**
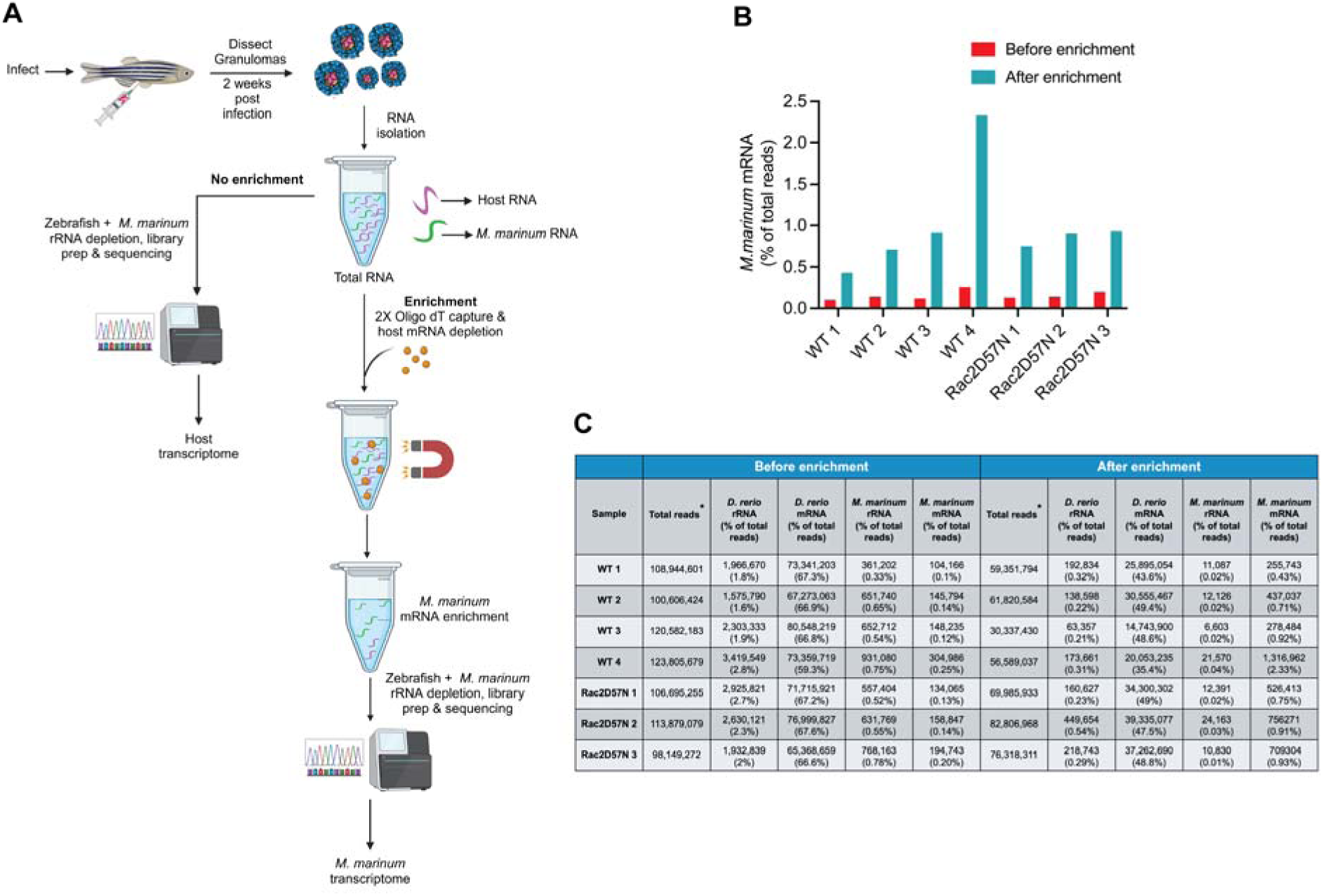
Dual RNA-Seq strategy for enriching mycobacterial transcripts from necrotic granulomas. **(A)** Diagram depicting the oligo(dT) bead-based host mRNA depletion strategy to enrich *M. marinum* transcripts from zebrafish necrotic granulomas. The figure was created with BioRender.com. **(B)** Bar plot showing enrichment of *M. marinum* transcripts in WT and Rac2D57N samples using the dual RNA-Seq approach. For each experiment, ∼250 granulomas from 8 WT/12 Rac2D57N fish were used to isolate total RNA. n = 4 independent experiments for the WT and 3 independent experiments for the Rac2D57N fish. **(C)** Table showing total reads (* filtered for low-quality reads) and the number of rRNA and mRNA reads for zebrafish and *M. marinum,* before and after enrichment.

### Neutrophils facilitate mycobacterial survival in granulomas via induction of the *devR* regulon

We first focused on gaining direct insights into the mycobacterial response to neutrophil deficiency by analyzing the *M. marinum* transcriptome from wild-type and Rac2D57N granulomas. In this study, we also included in vitro log-phase *M. marinum* culture as a secondary control. Principal component analysis of the bacterial transcripts revealed differential clustering of the wild-type, Rac2D57N, and *in vitro* broth culture samples, highlighting unique transcriptional responses associated with each condition (Fig. 4A). We identified a total of 270 *M. marinum* genes differentially expressed between wild-type and Rac2D57N samples, with 119 upregulated genes and 151 genes that are downregulated significantly in Rac2D57N granulomas (p_adj_ < 0.1) compared to wild-type granulomas (Fig. 4B and Table S4). Among these, 141 genes were found to have definitive homologs in *M. tuberculosis* (Table S4). Bacteria from the neutrophil-deficient Rac2D57N granulomas exhibited downregulation of genes encoding factors associated with intrinsic antibiotic resistance, including Mmpl5, a transmembrane transporter associated with Bedaquiline efflux (*54*); Rv2989, a transcriptional regulator involved in Isoniazid resistance by repressing *katG* (*55*); and Rv1217c and Rv1218c, which together forms a drug efflux pump conferring resistance to multiple antibiotics (*56*) (Fig. 4, C and F). Notably, the transcriptional activator *whiB7* was significantly downregulated in Rac2D57N samples, with similar trends observed for some of the genes in its regulon associated with antibiotic resistance, including *hflx, Rv1473, and Rv1258c* (Fig. 4, C and F). WhiB7 is a redox sensor that can be activated under redox stress (*57*). The differential regulation of the *nadABC* operon and *nadD* gene, which are involved in NAD^+^ biosynthesis and maintaining NAD^+^/NADH balance (*58*), in the bacteria from Rac2D57N granulomas could indicate a redox shift (Fig. 4, D and F). This shift may, in turn, influence *whib7* levels in mycobacteria within neutrophil-deficient granulomas, although the upstream signal driving this metabolic shift remains unclear.

**Fig. 4.**
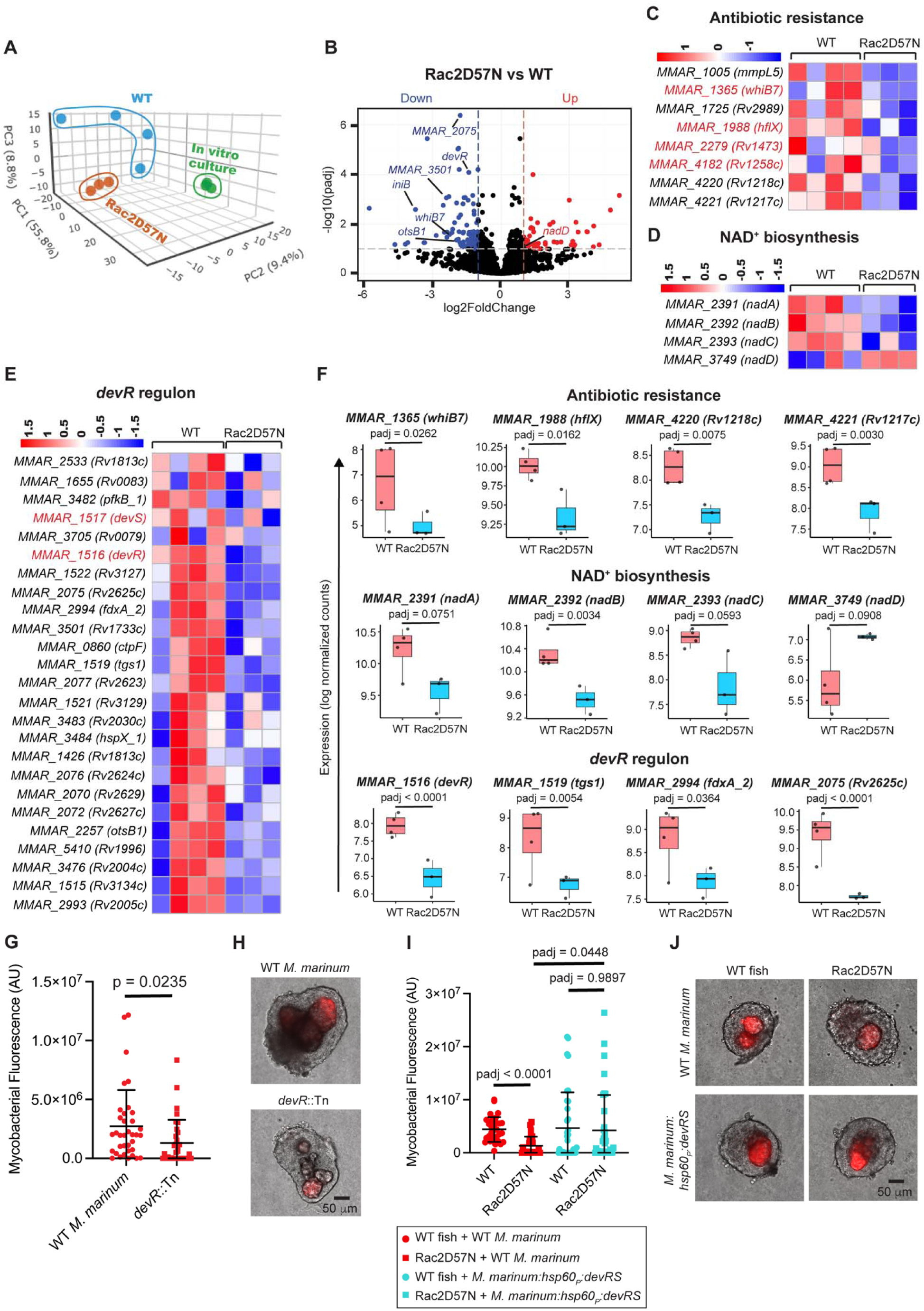
Neutrophils influence granuloma mycobacterial burden via *devR* regulon modulation. **(A)** Principal-component analysis (PCA) of the *M. marinum* transcriptome from wild-type and Rac2D57N granulomas and log phase in vitro broth culture. **(B)** Volcano plot showing differential expression of *M*. *marinum* genes in Rac2D57N vs WT granulomas. The horizontal dotted line indicates a p_adj_ threshold of 0.1, whereas the vertical dotted lines represent log2 fold change thresholds of −1 and 1 respectively. **(C, D, E)** Heatmaps showing relative expression levels for the genes associated with antibiotic resistance, NAD^+^ biosynthesis, and *devR* regulon in *M. marinum* from WT and Rac2D57N granulomas. **(F)** Box plots showing the median log normalized counts of representative transcripts associated with antibiotic resistance, NAD^+^ biosynthesis, and *devR* regulon in WT and Rac2D57N granulomas. Each data point represents the log normalized count of the respective transcript from an individual experiment. Benjamini-Hochberg (BH) correction was used to determine the adjusted p-values. **(G)** Bacterial burden in WT *M. marinum* and *devR* mutant-infected granulomas dissected from WT fish, represented as mean arbitrary fluorescence units (AU). For WT *M. marinum,* n = 36 granulomas; for *devR* mutant, n = 33 granulomas. Unpaired t-test with Welch’s correction. **(H)** Representative images showing WT *M. marinum* and *devR* mutant burden in the granulomas dissected from infected WT fish. **(I)** Granuloma bacterial burden in WT and Rac2D57N fish infected with either WT *M. marinum* (red data points) or *M. marinum* overexpressing *devR* & *devS* (cyan data points), represented as mean arbitrary fluorescence units (AU). For WT *M. marinum* infection: n = 32 granulomas from WT fish and 27 granulomas from Rac2D57N fish. For *M. marinum devRS* overexpressor infection: n = 35 granulomas from WT fish and 36 granulomas from Rac2D57N fish. Brown-Forsythe and Welch ANOVA tests, followed by Dunnett’s T3 multiple comparisons test, were used. **(J)** Representative images showing the burden of WT *M. marinum* and *M. marinum devRS* overexpressor in granulomas dissected from infected WT and Rac2D57N fish. **(G, I)** Granulomas were dissected from 14 dpi animals. Each data point represents mycobacterial fluorescence measured in a single granuloma, with granulomas obtained from 3 animals per group. Error bars indicate SD. **(H, J)** Scale bar, 50 µm.

A major highlight of the dataset was the downregulation of the mycobacterial two-component regulatory system *devR-devS* and its associated regulon in the neutrophil-defective granulomas (Fig. 4, E and F). DevS is a sensory membrane kinase activated in response to various conditions and signals present in the *M. tuberculosis* niche during intracellular infection or in necrotic granuloma lesions, including hypoxia, nitric oxide, and carbon monoxide (*59*). Upon activation, DevS phosphorylates the response regulator DevR, which then binds to upstream promoter regions and induces the expression of a set of genes called the *devR* regulon (*60*). This regulon comprises approximately 48 genes, some of which are associated with diverse functions, including alternate electron transport pathways (*fdxA*), nitrate metabolism (*narK2* and *narX*), and triglyceride synthesis (*<u>tgs1</u>*), contributing to *M. tuberculosis* persistence during chronic infection (*59, 60*). Of these 48 genes, we identified homologs for 25 genes in *M. marinum*, all of which were downregulated in neutrophil-deficient granulomas (Fig. 4E). In previous studies, *M. tuberculosis devR* mutants were reported to be attenuated in guinea pig, rabbit, and macaque infection models, underscoring the importance of this regulon in mycobacterial adaptation to necrotic granulomas (*61, 62*). To assess whether the the neutrophil-dependent *devR* downregulation could account for the effects of infection, we infected wild-type animals with an *M. marinum devR* mutant strain and compared its growth to that of the wild-type *M. marinum* within explant granulomas dissected 14 days post-infection. Consistent with the findings from mammalian models, we observed a significant reduction in the *devR* mutant burden in the granulomas compared to the wildtype strain, as determined through fluorescence measurements (Fig. 4, G, H and Fig. S6A). The downregulation of *devR* regulon (Fig. 4E), along with the reduced mycobacterial burden in neutrophil-deficient Rac2D57N granulomas (Fig. 2L), suggested that neutrophils may influence mycobacterial growth through transcriptional regulation of this regulon. In such a scenario, overexpressing this two-component system in *M. marinum* should restore the burden defect in the neutrophil-deficient granulomas. We indeed observed a comparable burden of *M. marinum* overexpressing the *devR*-*devS* via the *hsp60* constitutive promoter in both wild-type and Rac2D57N granulomas, indicating that the downregulation of the *devR* regulon is a contributing factor affecting mycobacterial growth in neutrophil deficient granulomas (Fig. 4, I, J and Fig. S6B, C).

### Neutrophil deficiency modulates inflammation and proliferation-associated transcriptional signatures

Analysis of the corresponding host transcriptome from wild-type and Rac2D57N granulomas revealed distinct transcriptional signatures between the two samples, as shown in the PCA plot (Fig. S7A). We identified 1,121 differentially expressed genes between the two samples, with Rac2D57N granulomas showing 482 upregulated and 639 downregulated genes (p_adj_ < 0.1) compared to wild-type granulomas (Fig. S7B and Table S5). To gain insights into the biological processes and pathways that are significantly altered in neutrophil deficient granulomas, we performed Gene Set Enrichment Analysis (GSEA) on the differentially expressed genes, which were ranked based on their fold change and compared them to predefined gene sets from Molecular Signatures Database (MSigDB). We found 10 gene sets from the Hallmark collection and 86 from the Curated collection (C2) in MSigDB that were significantly enriched in the Rac2D57N samples (p_adj_ < 0.05) (Fig. S7C, D, and Table S5).

The Hallmark data set revealed upregulation of gene sets associated with TNF-α signaling, IFN-γ, and inflammatory response pathways in Rac2D57N granulomas, suggesting disrupted immune homeostasis driven by neutrophil deficiency (Fig. S7C). TNF-α-signaling via the NF-κB pathway emerged as the most significantly enriched upregulated gene set in the hallmark collection (p_adj_ < 0.0005), with enhanced expression observed for *tnfa*, and associated transcriptional regulators including *maff, jun,* and *nfe2l2a* (*Nrf2*) (Fig. S7C, E, and H). Among these, NRF2 interacts with small Maf proteins like MafF and exerts cytoprotective functions by inducing the expression of antioxidant genes that shield against oxidative damage triggered by inflammation (*63*). It has also been shown to modulate the ability of alveolar macrophages to control *M. tuberculosis* growth during the early stages of infection (*64*). We observed upregulation of NRF2 target genes in Rac2D57N granulomas, indicating a compensatory response in which TNF-α-signaling-mediated oxidative stress induces *Nrf2* to maintain redox homeostasis (Fig. S7D, G, and J). A previous report demonstrated that neutrophils in macaque granulomas exhibit complex immunomodulatory properties, expressing both pro and anti-inflammatory cytokines (*65*). Additionally, there is evidence that neutrophils could exert immunoregulatory activity by inhibiting proinflammatory T-lymphocyte functions and their proliferation through mechanisms involving IL-10, arginase-I, and ROS (*66*). Our findings corroborate these observations, underscoring the role of neutrophils in regulating pro-inflammatory and oxidative stress response pathways in the granulomas.

Furthermore, we observed significant downregulation of genes associated with E2F transcription factors in neutrophil-deficient granulomas (Fig. S7C, F, and I). As key regulators of cell cycle progression, E2Fs can either promote or inhibit cell proliferation (*67*). The downregulation of genes associated with DNA replication and positive cell cycle regulation, such as *orc6, dscc1, cdc20,* and *cdca3* in this data set (Fig. S7I), suggested a possible decrease in the proliferation of specific cell populations within Rac2D57N granulomas. Previous scRNA-seq data has identified proliferative macrophage populations within granulomas (*36*). Our findings likely reflect a shift in cellular dynamics in neutrophil-deficient granulomas, although the specific cell types affected remain unclear.

### Dual RNA-Seq reveals mycobacterial transcriptional profiles unique to granulomas

Although our most specific focus was on the role of neutrophils, the development of the method to enrich bacterial transcripts allowed us to profile the totality of bacterial transcriptional programs induced *in vivo* in necrotic granulomas. We compared the mycobacterial transcripts from wildtype granulomas and in vitro log-phase broth culture, identifying 2,243 differentially expressed genes between the two samples (p_adj_ < 0.1) (Fig. 5A and Table S6). Of these, 1282 *M. marinum* genes (644 upregulated and 638 downregulated in wild-type granulomas) had clear homologs in *M. tuberculosis* (Table S6). These upregulated genes in the wild-type granulomas may represent both core pathways essential for mycobacterial survival across diverse host environments, such as the necrotic core and intracellular milieu, as well as pathways uniquely involved in adaptations to granuloma-specific conditions. To distinguish between these different gene categories and to identify granuloma-specific bacterial responses, we integrated our dataset with available *in vivo* mycobacterial transcriptional signatures obtained from *M. tuberculosis-*infected mouse lung macrophages (alveolar and interstitial) (*52*). We excluded 32 *M. marinum* genes with multiple *M. tuberculosis* homologs from our analysis to avoid ambiguities. Of the remaining 612 mycobacterial genes, 267 were commonly upregulated in both zebrafish wild-type granulomas and in vivo mouse macrophages (Fig. 5B and Table S7). This included genes associated with the synthesis and transport of the virulent cell wall lipid phthiocerol dimycocerosate (PDIM) and Kstr2 regulon involved in cholesterol catabolism, suggesting that these pathways may be essential for mycobacterial survival across granulomas and macrophages (Fig. 5, C, D and Table S7). Thus, we identified significant conservation of these upregulated genes in different model systems, irrespective of the differences in the host species and bacterial strain.

**Fig. 5.**
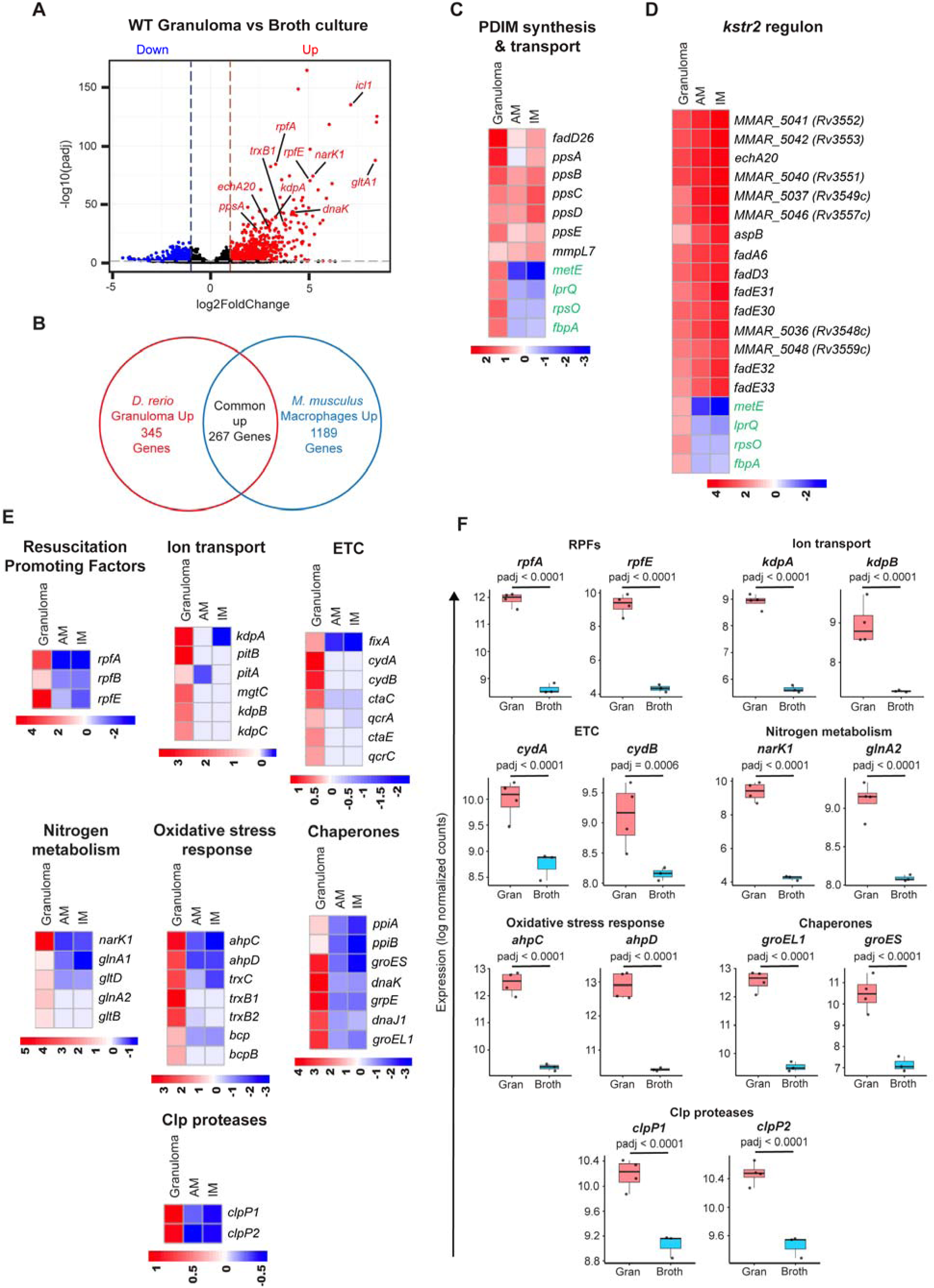
Dual RNA-seq identifies mycobacterial transcripts specifically enriched in necrotic wild-type granulomas. **(A)** Volcano plot showing differential expression of *M*. *marinum* genes in WT granulomas vs broth culture. The horizontal dotted line indicates a p_adj_ threshold of 0.1, whereas the vertical dotted lines represent log_2_ fold change thresholds of −1 and 1 respectively. **(B)** Venn diagram comparing upregulated mycobacterial genes from necrotic granulomas in zebrafish with previously published data from in vivo mouse macrophages. **(C, D)** Heatmaps showing mycobacterial genes commonly upregulated in zebrafish necrotic granulomas and in vivo mouse macrophages, related to PDIM synthesis & transport, and cholesterol catabolism. AM-alveolar macrophages, IM – interstitial macrophages. Four randomly selected genes specifically upregulated in necrotic granulomas (colored green) were included to improve scaling and avoid skewed representation. **(E)** Heatmaps showing mycobacterial genes specifically upregulated in zebrafish necrotic granulomas vs in vivo mouse macrophages, related to resuscitation promoting factors, ion transport, electron transport chain, nitrogen metabolism, oxidative stress response, chaperones, and Clp proteases. **(F)** Box plots showing the median log normalized counts of representative transcripts from **(E)** in WT granulomas and broth culture. Each data point represents the log normalized count of the respective transcript from an individual experiment. Benjamini-Hochberg (BH) correction was used to determine the adjusted p-values.

The remaining 345 genes, which did not overlap with the in vivo macrophage dataset and were uniquely upregulated in the necrotic granulomas *in vivo* (Fig. 5B and Table S8), were labeled as ‘granuloma-specific gene signatures.’ Analysis revealed that a subset of them belonged to well-annotated processes such as resuscitation from growth arrest, ion transport, electron transport chain, nitrogen metabolism, oxidative stress response, and unfolded protein response (chaperones and Clp proteases), suggestive of unique conditions that bacteria must endure specifically within the granulomatous environment (Fig. 5, E, F and Table S8). One proposed mechanism through which *M. tuberculosis* thrives in a hostile granuloma environment is its ability to enter a low metabolic, relatively inactive state under unfavorable conditions, which can be reversed upon the return of growth permissive conditions (*68*). *M. tuberculosis* encodes five genes named *rpfA-E,* which are believed to play a role in this process. Functional redundancy has been reported within this gene family as observed in mouse models during chronic infection, where single deletion mutants did not exhibit growth or resuscitation defects, but certain combinations of double or triple mutants did (*69*). The upregulation of multiple *rpf* genes (*rpfA, B,* and *E*), as observed in the granuloma dataset, highlights their potential importance for bacterial survival within granulomas (Fig. 5, E and F).

Increased expression levels of genes associated with ion transport, such as *kdpABC* (potassium transport), *pstA* and *pstB* (phosphate uptake), and *mgtC* (magnesium uptake) suggest that granulomas may impose specific ion limitations on mycobacteria (Fig. 5E). Notably, *M. tuberculosis* possesses two K^+^ uptake systems: one encoded by *kdpABC,* a high-affinity inducible system, and another, Trk, a constitutive system with low to moderate affinity. A previous report has established an essential role for Trk in mycobacterial survival within phagosomes, where K^+^ is not limiting (*70*). This raises the possibility that bacteria may switch between these two systems to maintain potassium homeostasis based on its availability under varied environmental conditions.

We observed a robust upregulation of the *cydAB* electron transport system, adapted for hypoxic environments (*71*), along with moderate upregulation of aerobic cytochrome *bc1-aa3-*related genes (*ctaC, ctaE, qcrA,* and *qcrC*) (Fig. 5E). This suggests that hypoxia is a dominant feature in granulomas, but some localized aerobic niches may exist that could sustain the function of the *bc1-aa3* pathway. We also inferred the metabolic status of mycobacteria in granulomas, particularly about fatty acid metabolism. Specific homologs involved in β-oxidation (*fadA, fadB*), lipopeptide biosynthesis, and fatty acid shuttling were identified (Fig. S8A and B). Notably, we observed upregulation of *echA6* but not its functional analog *fabH* in granulomas, suggesting a key role for *echA6* in fatty acid transfer and cell envelope lipid biosynthesis in this niche (Fig. S8B). Moreover, the requirement of genes like *fadA* for mycobacterial survival in necrotic granulomas in adult zebrafish (*72*) further strengthens their functional significance in this environment.

Additionally, the enrichment of a subset of nitrogen metabolism genes, as well as those related to oxidative stress response, protein misfolding (Fig. 5E), lipoprotein production, DNA biosynthesis, and transcriptional regulation (Fig. S8A and B), suggest that these gene sets may play a significant role in the adaptive responses of mycobacteria to the granulomatous environment. Overall, the comprehensive identification of the transcriptional modules unique to the necrotic granuloma microenvironment may serve as a useful resource.

### Evidence of evolutionary selection on transcripts enriched in necrotic granulomas

Granulomas are a key host-pathogen interface, and so bacterial genes required within them may be determinants of evolutionary success. We hypothesized that genes with transcripts enriched in necrotic granulomas are likely targets of natural selection during the long-term evolution of *M. tuberculosis*. We analyzed the *M. tuberculosis* homologs of the top 50 mycobacterial genes (ranked by their p_adj_ values) that we identified as specific to necrotic granulomas to test for evidence of evolutionary pressure and selection. We chose 69 fully sequenced *M. tuberculosis* and MTBC genomes across lineages and sub-lineages to first identify bacterial variants that were lineage-specific (Table S9). Specifically, we identified all variants across the 69 strains that had arisen once or multiple times and had been maintained across all strains in that lineage or sub-lineage without reverting, suggesting some adaptive advantage to the variation. In total, we identified 3,722 such mutational events. We then compared the frequency of these variants in the top 50 granuloma-specific transcripts to a simulated distribution of mutations occurring within the same set of genes under random conditions. We identified 39 non-synonymous lineage-specific variants in 23 of the 50 transcripts, representing a clear deviation (p<0.0001) compared to 1,000 simulations of random mutations in these genes (Fig. 6A and Table S9). These results suggest that genes responsive to necrotic granulomas have been more frequently mutated and then conserved during the long-term evolution of *M. tuberculosis*. Even synonymous variants were overrepresented in this gene list, although less significantly (p=0.001) (Fig. 6B and Table S9). Among the 23 genes, 9 showed multiple non-synonymous mutations, potentially indicating hotspots for adaptive variation (Table S9). These include *mmpl9,* encoding a transporter of an unknown substrate conferring resistance to oxidative stress (*73*), and *Rv1215c,* which codes for a conserved protein that interacts with EtfD, involved in fatty acid β-oxidation (*74*) (Fig. 6C). These findings suggest that transcripts that are specifically induced in this unique phase of the life cycle of pathogenic mycobacteria are targets of adaptation in the context of human infections.

**Fig. 6.**
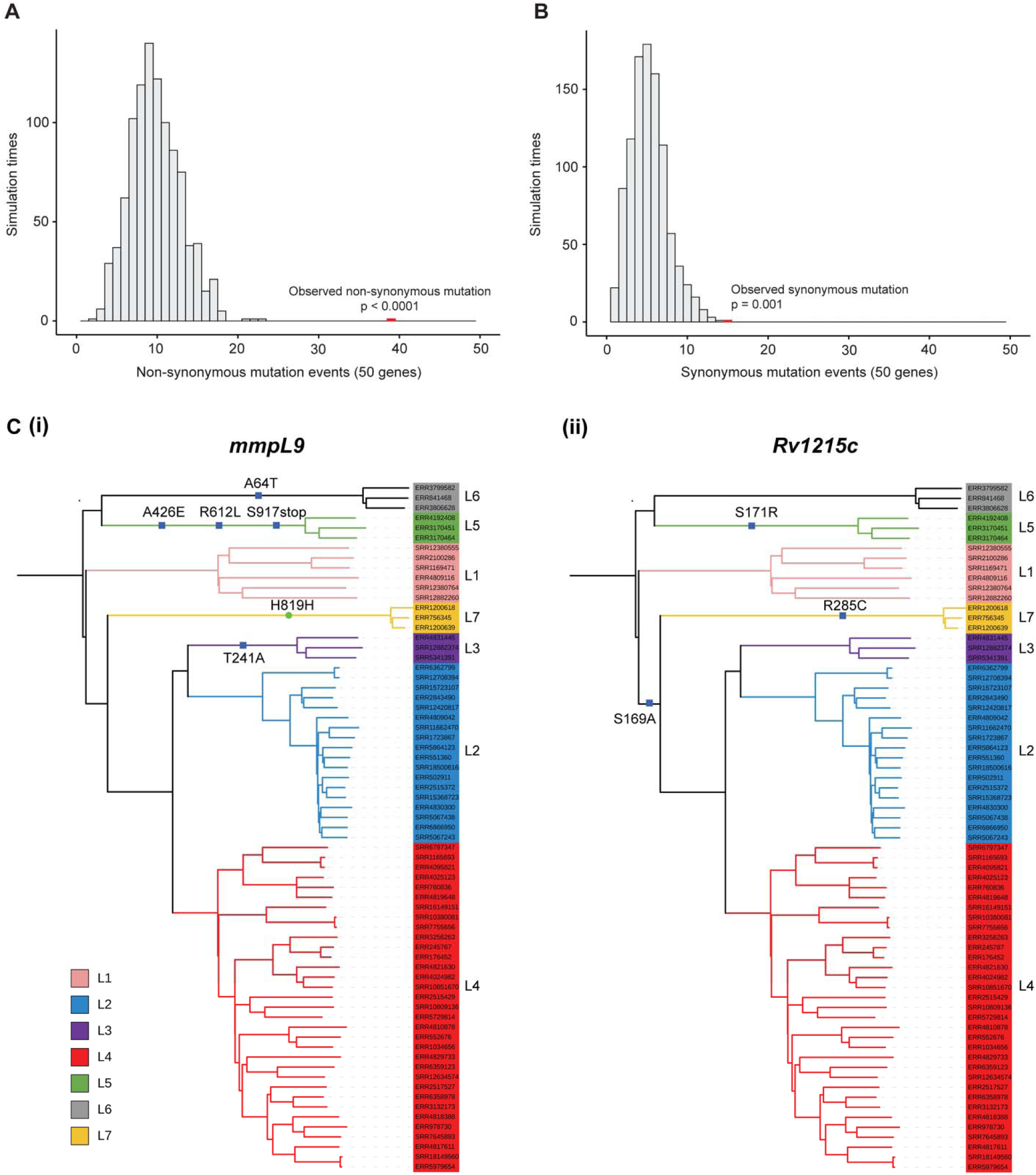
Lineage-specific adaptive variation in *M. tuberculosis* genes enriched in necrotic granulomas. (A, **B)** Distribution plots comparing the frequency of lineage-specific non-synonymous **(A)** or synonymous **(B)** mutations in the *M. tuberculosis* homologs of the top 50 granuloma-specific genes (red bars) to a simulated distribution of random mutations in this gene set (grey bars). n = 1,000 simulation events, a one-sided (right-tailed) permutation test was used to determine p-values. **(C)** Phylogenetic trees of 69 *M. tuberculosis* clinical isolates representing the seven primary lineages (L1-L7) and several major sub-lineages, including L4.1-L4.9 with mutational events for representative granuloma-specific genes **(i)** *mmpL9* and **(ii)** *Rv1215c* highlighted. Blue squares represent non-synonymous mutations, and the green circle represents a synonymous mutation. Branch colors indicate the inferred lineages.

## Discussion

Here, we employed a multifaceted approach combining high-resolution longitudinal imaging, explant culture techniques, pharmacological and genetic intervention strategies, and novel dual host-pathogen transcriptomics to investigate host-mycobacterial dynamics as well as mycobacterial persistence within highly organized necrotic granulomas. We first focused specifically on neutrophils, whose role in TB disease pathogenesis is complex and challenging to study owing to their short life span and the limitations of existing cell lines in accurately mimicking their behavior ex vivo (*75*). To address these challenges, we took advantage of a mycobacterial granuloma explant system, which recapitulates important features of human TB granulomas, such as diverse cellular compositions, inflammatory heterogeneity, epithelioid morphology, and a well-defined necrotic core (*4*), to gain deeper insights into neutrophil functions within these complex structures.

Live imaging of the explant granulomas, coupled with scRNA-seq analysis revealed phenotypic, functional, and transcriptional diversity among neutrophils shaped by distinct microenvironments within the granulomas. Neutrophils are known to exhibit functional heterogeneity both in the basal state and during inflammatory and disease conditions, such as cancer. In particular, their heterogeneity in cancer has been extensively studied, leading to the characterization of anti-tumorigenic N1 neutrophils, pro-tumorigenic N2 neutrophils, granulocytic myeloid-derived suppressor cells (G-MDSC), and low-density neutrophils (LDN) with T-cell suppressive functions (*29*). Taking a cue from cancer studies, neutrophil heterogeneity in the context of TB has been previously explored based on neutrophil inflammatory and maturation status, leading to classifications into MDSC and LDN-like cell types with immunosuppressive functions, as well as steady-state neutrophils (SSNs), which are relatively more mature (*66*). However, their relevance to necrotic granulomas which exhibit a complex cellular and inflammatory milieu, remains unclear. We identified a subset of the granuloma neutrophil population enriched with transcripts associated with granulocyte maturation markers and pro-inflammatory cytokine. This suggests that functional divergence in the granuloma neutrophil population may exist based on their maturity and cytokine profile.

Epithelial cell layers are characterized by their basal-apical polarization and tightly interdigitated organization, facilitated by desmosomes, adherens junctions, and tight junctions. Upon recognizing inflammatory stimuli, neutrophils interact with these epithelial layers via multiple adhesion molecules, allowing them to squeeze and traverse through the epithelial barrier (*76*). In previous work, we identified that granuloma macrophages, especially those surrounding the necrotic core, undergo epithelial reprogramming analogous to mesenchymal-epithelial transitions, resulting in the expression of canonical epithelial markers such as E-cadherin, plakoglobin, and α-catenin (*50*). The highly dynamic, elongated neutrophils observed in type I and type II granulomas were predominantly seen patrolling and traversing these epithelialized macrophage layers surrounding the necrotic core. This suggests that their interactions with these epithelioid macrophages may shape neutrophil morphology and behavior while they transmigrate through these cell layers. Such interactions could transiently compromise the barrier, possibly altering the signaling events occurring between the epithelialized macrophages For example, elastase released by migrating neutrophils has been shown to cleave E-cadherin, thereby activating proliferative signals in epithelial cells (*77*). In line with this, we observed a significant downregulation of gene signatures associated with cell proliferation in neutrophil-deficient granulomas. Thus, interactions between neutrophils and epithelioid macrophages observed in specific granuloma compartments may mutually influence each other, shaping their cellular behavior and signaling events.

Pharmacological and genetic interventions revealed a crucial role for PI3Kγ/δ and the Rac2 signaling axis in neutrophil-mediated support of mycobacterial survival in granulomas. The reduction in mycobacterial burden observed following the chemical targeting of this signaling pathway with the FDA-approved drug Duvelisib serves as proof of principle, opening avenues for exploring host-directed strategies that target neutrophils as a complementary approach to tackling TB infection. In recent years, significant interest has grown in developing host-directed therapies for TB, which offer a way to overcome the limitations posed by the emergence of *M. tuberculosis* strains resistant to current drugs (*78*). While targeting host factors in neutrophils is a promising approach, caution is needed to avoid disturbing their core antimicrobial functions, which may lead to unintended consequences such as increased susceptibility to secondary infections. A more refined approach distinguishing pathogenic from protective neutrophil populations and localized inhibition of detrimental neutrophil functions may ultimately offer safer and more effective therapeutic interventions.

While host-centric processes such as NETosis and the metabolic profile of neutrophils have been previously linked to supporting *M. tuberculosis* survival (*14–18*), their direct influence on mycobacterial physiology and adaptation, eventually affecting bacterial growth, remains unexplored. The dual RNA-seq strategy we developed in this study allowed us to simultaneously capture both host and mycobacterial transcriptional changes in neutrophil-deficient granulomas, offering a deeper understanding of how neutrophils affect mycobacterial responses and contribute to their survival in the granulomatous niche. In parallel, the ability to access the bacterial transcriptome from the granulomas has enabled us to identify genes specifically associated with mycobacterial adaptation to the necrotic lesions. This complements previous dual RNA seq approaches, which were primarily focused on identifying mycobacterial genes that support adaptations to intracellular niches (*51, 52*). Technically, our approach is relatively simple and captures transcripts almost immediately after granuloma harvest via direct lysis of granuloma samples in buffer. Additionally, the use of an oligo(dT) bead depletion strategy to remove poly(A)-tail containing host mRNAs allows for relatively unbiased representation of the mycobacterial transcriptome. Further, by including around 400 granulomas per sample for each condition, the strategy likely encompasses heterogeneous granuloma subtypes resulting from variations in cellular composition and microenvironment.

Through dual RNA-seq, we identified a role for neutrophils in supporting mycobacterial survival in granulomas via upregulation of DevR-DevS-mediated adaption mechanisms. This two-component system in mycobacteria has been previously shown to be activated under stress conditions typical of the mycobacterial niche, such as hypoxia, nitric oxide, and carbon monoxide, as well as during intracellular survival in macrophages (*52, 59*). Our study linking neutrophils to the activation of the DevR-DevS in necrotic granulomas introduces a new dimension to the understanding of its regulation. This suggests that mycobacteria can respond to some functions of neutrophils through this two-component system, promoting their persistent survival in granulomas. Currently, it remains unclear whether this neutrophil-mediated regulation is direct or occurs indirectly through its interactions with other immune cells. However, our observation of direct interactions between dying neutrophils and mycobacteria in specific granuloma microenvironments suggests that this regulation might be direct, possibly via the release of yet-to-be-characterized signals from neutrophils. With DevS being a gas sensor, one possible candidate could be the nitric oxide produced by neutrophils at higher levels, particularly during hypoxic conditions that prevail in granulomas (*19*). In such a scenario, the harmful effects of nitric oxide could be mitigated by the virulence mechanisms that exist in mycobacteria (*79*), enabling it to safely sense this signal. However, these propositions require further evaluation. These findings, along with the previous studies (*15, 18*), present a complex picture of neutrophil-mediated mycobacterial growth regulation during TB infection, where host processes such as NETosis and bacterial adaptation mechanisms through the DevR-DevS may synergistically contribute to mycobacterial survival.

Granulomas are believed to provide a complex survival environment for *M. tuberculosis,* and our understanding of how mycobacteria dynamically regulate their genes and adapt during chronic infection in these structures remains an underexplored topic. We identified granuloma-specific mycobacterial transcripts that are reflective of their unique metabolic state, as well as the nutritional and stress environments prevailing in the necrotic core. While previous works primarily focused on metal homeostasis, including iron, copper, and zinc in *in vivo* macrophages (*80*), our identification of granuloma-specific gene sets associated with potassium, magnesium, and phosphate transport suggests that *M. tuberculosis* may require a diverse set of nutrients and cues for its extracellular persistence in granulomas that are not amenable to study in other systems. The transport systems encoded by these genes are relatively unexplored, and their characterization could significantly expand our understanding of nutrient acquisition strategies of *M. tuberculosis* in granulomas. The functions of several genes identified in this study, like those encoding resuscitation-promoting factors and cytochrome *bd* electron transport system, have been linked to granuloma-related processes primarily through annotation and indirect evidence from prior in vitro, ex vivo, and mouse studies (*68, 69, 71*). The robust upregulation of oxidative stress response genes and chaperones suggests that mycobacteria may experience significantly higher stress levels in granulomas as opposed to intracellular environments.

This study highlights potential pathways that can be leveraged to disrupt mycobacterial adaptation in necrotic lesions and improve treatment outcomes. Many first and second-line TB drugs effectively target actively growing bacteria, but they become less effective when *M. tuberculosis* is subjected to conditions similar to those that prevail in the granulomas. They adopt a drug-tolerant state with possible alterations in their metabolic and stress profiles, a phenomenon that could likely contribute to the prolonged treatment regimen required for TB (*81*). Identification of genes that may be crucial for mycobacterial survival in granulomas may provide insights into potential vulnerabilities such as metabolic adaptations, stress response mechanisms, or persistence-associated pathways that could be targeted to shorten treatment duration and enhance therapeutic efficacy.

We interrogated a diverse set of *M. tuberculosis* strains and identified lineage-specific adaptations in the granuloma-associated genes. We found enrichment of non-synonymous mutations that were then retained throughout diverse lineages and sublineages, suggesting that these genes are under specific selective evolutionary pressures. Mycobacterial survival and adaptation to the granuloma microenvironment is likely a common feature of bacterial evolutionary success across lineages and throughout diverse human populations.

## Materials and Methods

### Animal handling and maintenance

Zebrafish husbandry and experiments were performed in compliance with the National Institutes of Health standards for the care and use of animals and approved by the Duke University Animal Care and Use Committee (protocol A049–23-03). Adult zebrafish were housed in 3 or 6 L tanks on a 14-hour light/10-hour dark cycle and maintained under the following water conditions: temperature: 28°C; conductivity: 600-700 μS (maintained with Instant Ocean Sea Salt, (#SS15-10)); pH: 7.0-7.3 (buffered by sodium bicarbonate; Arm & Hammer Pure Baking Soda (#426292)). Animals were fed twice daily – once with dry food and once with *Artemia*.

Larval zebrafish were maintained at 28.5°C in 100 mm Petri dishes in sterile E3 media (5 mM NaCl, 178 μM KCl, 328 μM CaCl_2_, 400 μM MgCl_2_) with a density of around 60 larvae per dish. For imaging, pigmentation was arrested by adding 45 μg/ml 1-phenyl-2-thiourea (PTU, Sigma-Aldrich cat# P7629) at 1day postfertilization (dpf).

### Zebrafish wild-type and transgenic lines

All zebrafish strains were in the *AB wild-type background. The transgenic lines Tg(*lyz:egfp*)*^nz117^*, and Tg(*irg1:tdTomato*) *^xt40^* have been previously described (*26, 33*). Tg(*mpx:mcherry*-2A-*rac2*D57N) (Rac2D57N fish) was a gift from A. Huttenlocher (*49*). Due to the possible variations in the transgene copy numbers in the Rac2D57N line, we observed both brighter and dimmer progenies after spawning. These larvae were sorted based on fluorescence intensity, and the brighter ones were selected for further maintenance.

### Bacterial strains and culture conditions

The bacterial strains used in this study were derived from the wild-type *M. marinum* M strain. *M. marinum* containing *msp12:tdTomato* and *msp12:mCerulean* were described previously (*83, 84*), and they were grown in Hygromycin B-containing media.

The transposon mutant *devR*::Tn was identified from a sequenced library of *M. marinum* transposon mutants, kindly provided by C. Cosma and L. Ramakrishnan. The transposon insertion site was confirmed by PCR and sequencing, as described previously (*85*). The following primers were used for confirmation: *devR*::Tn check F: 5’-GAGTGGTCCGTTGGTAAC-3’(anneals upstream of *devR*), *devR*::Tn check R: 5’-ATGTGGTTCGTCGGTCAT-3’ (anneals downstream of *devR*), and TnMarR3: 5’-ACAACAAAGCTCTCACCAAC-3’ (anneals to the transposon). The transposon insertion cassette (TnMar) in the mutant confers resistance to Hygromycin B. A Fluorescent *devR*::Tn strain was then generated by electroporating the plasmid *msp12:mCerulean*-KanR (a gift from L. Ramakrishnan), which drives the expression of the mCerulean fluorescent protein and confers resistance to Kanamycin.

To generate the *devR*-*devS* overexpressor strain, the *devR-devS* open reading frames, along with the intergenic region in *M. marinum*, were amplified and cloned next to the constitutive *hsp60* promoter in pMCHP-KanR (*85*) by replacing the pre-existing *pcaA* in that plasmid. The following primers were used for this step: pMCHP F: 5’-CAGCTGATATCCCCTATAGTG-3’, pMCHP R: 5’-CATTGCGAAGTGATTCCTC-3’ (To linearize the pMCHP-KanR), MarDevRS F: 5’-AATCACTTCGCAATGGCCAAGACAATCGCGGCTTGGTAACGGTCTTCTTGG-3’, MarDevRS R: 5’-AGGGGATATCAGCTGTGGCCCGAACAACTATTG-3’ (for *devR-devS* open reading frame amplification). The linearized plasmid and the insert were then ligated together using In-Fusion (TaKaRa) cloning. The *hsp60* promoter*-devR-devS* region was then amplified from this plasmid using hsp60 F (5’-TCTCATCAACCGTGGAAATCTAGAGGTGACCACAAC-3’) and MarDevS R2 (5’-CAGAAAGTGAGGGAGTGTTGGCTAGCTGATCAC-3’) primers. The amplified fragment was subsequently cloned into the *msp12:cerulean* plasmid using In-Fusion cloning. The plasmid was linearized using the primers msp12 mcerulean F (5’-CTCCCTCACTTTCTGGC-3’) and msp12 mcerulean R (5’-CCACGGTTGATGAGAGC-3’). The resulting plasmid, pMCHDevRS, which constitutively drives the expression of *devR-devS* along with the mCerulean fluorescent protein, was electroporated into the wild-type *M. marinum* strain and selected using Hygromycin B.

The bacterial cultures were grown to an OD_600_ of ∼ 1.0 in 7H9 complete media supplemented with 10% OADC (50 g/L BSA, 0.5% oleic acid, 20 g/L dextrose, 8.5 g/L NaCl) and 0.05% Tween-80. As appropriate, either 50 ug/ml Hygromycin B or 20 ug/ml Kanamycin was added. Single-cell aliquots of *M. marinum* cultures were prepared for adult and larval fish infections as described previously (*46*), by passing them through a 27G needle and a 5μm filter. After bacterial enumeration, the aliquots were stored at −80°C.

### Adult zebrafish infections

Adult zebrafish infections were performed as previously described (*36*). Fish were anesthetized with 0.016% tricaine and injected intraperitoneally with ∼ 350 fluorescent *M. marinum* strain. After injection, they were allowed to recover in clean fish system water and maintained at 28.5 °C under a 14-hour light/10-hour dark cycle for 14 days in an incubator. Water was changed every day, and the animals were fed daily. They were monitored regularly and euthanized if they showed signs of severe distress or appeared to be dying.

### Granuloma explant culture (Myco-GEM)

Granulomas were harvested from adult zebrafish 14 days post-infection (dpi) with *M. marinum* and cultured using the Myco-GEM technique as described previously (*4*). Briefly, infected adult fish were euthanized, and granulomas were dissected from the body cavity in L-15 medium (Gibco #21083–027). These granulomas were washed three times in L-15 medium and transferred to an optical-bottom 96-well plate (GBO #55090) covered with 40 µl of 5 µg/ml Cultrex® Basement Membrane Extract, (R&D systems #3432-005-01), in L-15. The granulomas were then embedded in a top medium containing 5% FBS (Sigma #2442) and 1 µg/ml Basement Membrane Extract in L-15 and subsequently imaged.

### Confocal imaging and image analysis

Time-lapse images of granuloma explants were acquired with Crest X-Light V2 spinning disk confocal system (CrestOptics) equipped with LDI-NIR laser diode illuminator (89 North, USA), ORCA-Flash4.0 V3 Digital CMOS camera (Hamamatsu), and Zeiss Observer Z1 inverted microscope. 20X/0.5 EC “Plan-Neofluar” (Zeiss) lens was used to capture the images. Z stacks were taken at 5 µm and maximum intensity projections were made using ImageJ software. Neutrophil circularity was measured using the ‘Analyze particles’ menu command in ImageJ. Briefly, a threshold range was set to differentiate neutrophils from the background, and the neutrophils were randomly selected using the Wand tool for further measurement. Neutrophil tracking and velocity measurements were carried out using the MTrackJ plugin in ImageJ. Neutrophils were randomly selected for the velocity measurements. Neutrophil death was quantified visually by subtracting the number of fluorescent neutrophils in type I explant granulomas/ type II ENR 13 hours post-dissection from the initial neutrophil numbers observed immediately after dissection. The percentage of neutrophil death was subsequently calculated based on these measurements. The near-complete compartmentalization of elongated and rounded neutrophils within the granuloma microenvironment facilitated the quantification of their survival with less difficulty.

### Zebrafish sectioning

Infected zebrafish were euthanized and fixed in CLARITY hydrogel solution (4% paraformaldehyde, 4% acrylamide, 0.05% bis acrylamide, 0.0025 g/ml VA-044) for 2 days and processed further as described previously (*36*). The hydrogel containing fixed animals was polymerized for 3 hours at 37°C, and excess hydrogel was removed from the tissues. The tissues were then sequentially incubated in 10% sucrose/1x PBS, 20% sucrose/PBS, and 30% sucrose/PBS for 1 day each at 4°C, followed by their incubation in a 50/50 mix of Neg-50 (Epredia #22-110-617) and 30% sucrose/PBS at room temperature for an hour. The tissues were finally frozen in pure Neg-50 and sectioned at 20 µm using a cryostat.

For imaging, slides containing these sections were mounted in DAPI Fluoromount-G (SouthernBiotech #0100-20). The sections were imaged with the Crest X-Light V2 spinning disk confocal system described above using a 63X/1.15 LD “C-Apochromat” W Corr (Zeiss) water immersion lens. Z stacks were acquired at 2 µm intervals, and maximum intensity projections were generated using the ImageJ software.

### Single-cell RNA-seq analysis of granuloma neutrophils

Single-cell RNA-seq (ScRNA-seq) analysis of granuloma-associated neutrophils was performed by extracting and analyzing the neutrophil population from the previously published zebrafish granuloma ScRNA-seq dataset (GSE161712) (*36*). The neutrophil population was subclustered using the R package Seurat v3.4 (*86*). The statistical significance of the PCA scores was assessed using the JackStraw() and ScoreJackStraw() functions, and four significant principal components were selected for subsequent analyses. The FindNeighbors() and FindClusters() functions identified three neutrophil subclusters, with their corresponding markers determined using the FindConservedMarkers() and FindAllMarkers() functions. UMAP plots were used to visualize these neutrophil subclusters and the differential expression of relevant subcluster-specific genes. Gene Ontology (GO) analysis of differentially expressed genes within each subcluster (p_adj_ < 0.1) was carried out using the gprofiler2 package (*87*).

### Duvelisib treatment of explant granulomas

A 10 mM stock of Duvelisib (Selleckchem #S7028) was made in DMSO and stored at −80°C. Fresh dilutions of the drug were made for each experiment so that the granuloma explants were treated with the final concentration of 5 µM Duvelisib in 1% DMSO in top media. Control granulomas were treated with 1% DMSO in top media. Neutrophil circularity and velocity were measured 5 hours post-treatment with Duvelisib or vehicle.

### Quantification of mycobacterial burden in explant granulomas

For measuring *M. marinum* burden in Duvelisib-treated and control explant granulomas, they were imaged on day 0 and day 4 post-treatment using epifluorescence microscopy (inverted Zeiss Observer Z1 microscope) with a 20X objective. Exposure time was kept constant throughout imaging. Images were analyzed using ImageJ, setting a constant threshold above the background. *M. marinum* burden was expressed as a function of its fluorescence, calculated by multiplying the mean fluorescence intensity by the fluorescence area in granulomas, as described earlier (*46*). No significant difference in *M. marinum* burden was observed between the control and Duvelisib treatment groups on day 0.

Due to possible variations in *mpx* promoter activity, we observed differences in Rac2D57N levels among circulating neutrophils, as reflected in their fluorescence intensity across adult fish, which became evident after dissection. Therefore, for bacterial burden measurements and other experiments in Rac2D57N explant granulomas, we selected only those fish with brighter neutrophils.

### In vitro growth kinetics

Wild-type *M. marinum* grown to late exponential phase was diluted to an OD_600_ of ∼ 0.2 and cultured in 7H9 complete media with 5 µM Duvelisib or 1% DMSO at 33°C in a shaker set to 150 RPM. Growth curves were generated by plotting OD_600_ measurements against time.

### Mycobacterial colony-forming unit (CFU) determination in adult fish

Adult fish at 14 days post-infection with *M. marinum* strains were euthanized using a Tricaine overdose. Each fish was placed in a 2 ml screwcap tube prefilled with 2.8mm stainless steel beads (Sigma #Z763829-50EA), one 6.5 mm ceramic bead (Omni #19-682), and 1 ml of 7H9 containing 0.05% Tween-80 and 50 ug/ml Hygromycin B. The samples were homogenized in a bead mill for two cycles of 15 seconds each and then serially diluted in 7H9 containing Tween-80 and Hygromycin B. The serial dilutions were plated on 7H10 agar supplemented with 10% OADC, 50 ug/ml Hygromycin B, 25 ug/ml Polymyxin B, and 10 ug/ml Amphotericin B. The plates were incubated at 30°C for ∼10 days, and the colonies were enumerated.

### Hindbrain ventricle (HBV) infection in larvae

Hindbrain ventricle infection of Tg(*lyz:egfp*) and double transgenic Tg(*lyz:egfp*);Tg(*mpx:mcherry*-2A-*rac2*D57N) larvae with wildtype *M. marinum* containing *msp12:tdTomato* was performed as described previously (*20*). At 2 dpf, each larva was infected with ∼100 fluorescent *M. marinum* in the hindbrain ventricle using the Vacuum-Assisted MicroProbe (VAMP), alongside PBS-injected (Mock) controls. The number of neutrophils recruited to the HBV was enumerated on day 3 post-infection from confocal images.

### Total RNA extraction and Dual RNA-seq of granulomas

For each biological replicate, around 400 granulomas were dissected from 8-12 wild-type or 14-16 Rac2D57N adult zebrafish infected with wild-type *M. marinum* at 14 dpi and transferred to 2 ml O-ring tubes in L-15 medium. After removing the medium, granulomas were washed with 1 ml of 1X PBS and resuspended in RLT plus buffer supplied with the RNeasy Plus Mini kit (QIAGEN #74134). The suspended granulomas were then lysed using 0.7 mm zirconia beads (BioSpec Products #11079107zx) in a BeadBug^TM^3 homogenizer (Benchmark Scientific #D1030) at 4000 RPM for 35 s, and this process was repeated four more times. Total RNA was then isolated using the RNeasy Plus Mini kit by following the manufacturer’s protocol.

The Sequencing and Genomics Technologies Core Facility at Duke University performed mRNA-sequencing. Total RNA was split into two portions: one was processed without bacterial mRNA enrichment to get the host mRNA reads, while the other was subjected to *M. marinum* mRNA enrichment by following a two-step approach. First, zebrafish mRNA was removed from total RNA using two oligo dT captures (Kapa mRNA HyperPrep Kit #KK8581). The remaining RNA in the supernatant was then reverse transcribed into cDNA and processed into sequencing libraries using the Illumina Total RNA-seq with Ribo-Zero Plus Microbiome kit (Illumina #20072063), with additional custom rRNA depletion probes for zebrafish and *M. marinum* (Table S3). Total RNA from in vitro *M. marinum* log phase cultures were treated similarly. Libraries from individual samples were pooled at equimolar concentration and sequenced on the Illumina NovaSeq 6000 S Prime flow cell to generate 150 bp paired-end reads.

### Dual RNA-seq data processing and analysis

Raw sequencing reads were quality-checked using FastQC (v. 0.11.9), and low-quality reads were filtered, and Illumina adapters were trimmed using fastp (v. 0.22.0) (*88*). Bowtie2 (*89*) was used to assess rRNA content in the raw reads with custom-built zebrafish and *M. marinum* rRNA indexes. Filtered and trimmed reads were pseudoaligned individually to the zebrafish and *M. marinum* transcriptome indexes built from their corresponding transcriptomes (GRCz11 and GCA_000018345.1 (ASM1834v1), respectively) using Kallisto (v. 0.46.1) (*90*), which quantified transcripts abundances.

Transcript-level abundance estimates were then imported separately for zebrafish and *M. marinum*, aggregated to the gene level, and normalized based on transcript abundance and length using the tximport () function in R. Differential gene expression analysis was performed using the DESeq2 package (v. 1.44.0) (*91*), and heatmaps for differentially expressed genes were generated using pheatmap (v. 1.0.12).

For Gene Set Enrichment Analysis (GSEA) of differentially expressed host transcripts, log fold change (LFC) shrinkage was performed using apeglm (v. 1.26.1) to stabilize fold change estimates for lowly expressed genes and improve gene prioritization (*92*). Genes were then ranked based on their LFC values, and GSEA was conducted with the clusterProfiler (v. 4.12.0) package (*93*) using gene sets from the Molecular Signatures Database (MSigDB) (https://www.gsea-msigdb.org/gsea/msigdb). Enrichment plots were generated using enrichplot (v. 1.24.0).

One of the Rac2D57N samples was identified as an outlier in the PCA analysis of the *M. marinum* transcriptome and had 2-3 times fewer bacterial reads than other samples before enrichment, suggesting loss of bacterial RNA during processing. Therefore, this sample was excluded from our analysis.

### Identification of granuloma-specific mycobacterial gene signatures

Granuloma-specific mycobacterial gene signatures were identified by comparing the *M. tuberculosis* homologs of 612 *M. marinum* genes that were significantly overexpressed in wild-type necrotic granulomas, relative to *M. marinum* log-phase broth cultures, with previously published in vivo mycobacterial transcriptional signatures (*52*). These in vivo signatures consist of 1,456 *M. tuberculosis* genes that are significantly upregulated during intracellular survival in mouse lung alveolar and interstitial macrophages. Comparisons were made using the tidyverse (v. 2.0.0) package in R. Of these, 267 genes were found to be commonly upregulated in both zebrafish wild-type granulomas and in vivo mouse macrophages. The remaining 345 genes, which did not overlap with the in vivo macrophage dataset, were identified as granuloma-specific gene signatures.

### Identification of lineage-specific adaptive variations in granuloma-associated *M. tuberculosis* genes

Fully sequenced genomes of 69 *M. tuberculosis* clinical isolates, representing the seven primary lineages (L1-L7) and several major sublineages, including L4.1-L4.9 and *Mycobacterium bovis* (Table S9) (*94*), were analyzed. Phylogenetic analysis was conducted following our previous method (*94*). Ancestral sequences for each node in the phylogeny were inferred using IQ-TREE (v. 2.2.2.7) with the GTR nucleotide substitution model. Lineage-defining mutations at each node were identified by comparing the ancestral sequence of each node to that of its parent. In total, 2,413 nonsynonymous mutations and 1,309 synonymous mutations were identified.

From these, we selected mutations within *M. tuberculosis* homologs of the top 50 granuloma-specific genes (ranked by p_adj_ value) (Table S8), identifying 39 nonsynonymous and 15 synonymous mutations.

To assess whether these mutations indicate positive selection, we simulated the number of mutations that would occur in these top 50 genes under random conditions. All *M. tuberculosis* lineage-defining mutations were randomly distributed across approximately 4,111 genes, with mutation probabilities adjusted for gene length for accuracy. This simulation was repeated 1,000 times, and the resulting distribution of mutations in the top 50 genes under random conditions was used as a baseline for comparison.

A one-sided (right-tailed) permutation test was used to determine statistical significance. p-values were calculated based on the cumulative probability of the simulated distribution. Since the observed values lie on the right side of the distribution, we calculated the proportion of simulated values greater than or equal to the observed value to determine the p-values.

### Statistics

Statistical analyses for scRNA-seq and Dual RNA-Seq were performed using the packages mentioned in the respective sections. For all other experiments, statistical analysis was conducted using Prism 10 (GraphPad Software).

## Supporting information

Combined Supplementary Figures

## Data availability

Raw sequencing reads and count matrices for Dual RNA-Seq are available in GEO under accession number GSE289727.

## Acknowledgments

We thank A. Huttenlocher for the Rac2D57N fish line, C. Cosma and L. Ramakrishnan for the *M. marinum* transposon mutant library, and M. Cronan, C.M. Smith and members of the Smith and Tobin laboratories for helpful discussions and comments on the manuscript. This work was funded by National Institutes of Health grants AI125517 and AI130236 (D.M.T.). The funders had no role in study design, data collection and analysis, decision to publish, or preparation of the manuscript.

## Author contributions

G.V. and D.M.T. conceived and designed the study. G.V. performed and analyzed all zebrafish experiments and processed and analyzed the Dual RNA-Seq data. E.J.H. performed single-cell RNA-seq analysis. G.A. carried out *M. marinum* mRNA enrichment, cDNA library preparation, and sequencing for the Dual RNA-Seq experiment under the supervision of D.S.-L. M.G. and Q.L. identified lineage-specific adaptive variations and performed phylogenetic analysis with supervision from Q.L. A.M.X.-M. generated *M. marinum devR*::Tn fluorescent and *devR*-*devS* overexpressor strains and prepared single-cell *M. marinum* cultures. G.V. and D.M.T. wrote the manuscript with substantial contributions from all authors.

## Supplementary figure legends

**Fig. S1. Neutrophils migrate between sub-compartments of a type II granuloma.** Movement of two neutrophils tracked over 220 mins (red and white arrows) from a region near the necrotic core to the ENR (white dashed box) of a type II granuloma. Scale bar, 50 µm.

**Fig. S2. Neutrophil death in type II granulomas is independent of the cell infection status.** Magnified images of the extra–necrotic region (ENR) of a type II granuloma shown in **Fig. 1G** at different time points. The red arrow indicates an uninfected neutrophil; the white arrow indicates an infected neutrophil. Scale bar, 25 µm.

**Fig. S3. Granuloma microenvironment governs neutrophil macrophage interactions. (A)** Fluorescent time-lapse images showing macrophage (red) and neutrophil (green) viability in *M. marinum* (magenta) - infected type I and ENR (white box) of type II granulomas observed over 780 mins. Images are representative of granulomas from three independent experiments, n = 9 animals, scale bar, 50 µm. **(B)** Magnified images of the ENR (white boxes in panel A) at different time points, showing interactions between a dying neutrophil (red arrow) and an infected macrophage (white arrow). The yellow arrow indicates *M. marinum* released from the infected macrophage after its interaction with the neutrophil. Scale bar, 20 µm. Fluorescent images are 100 µm maximum projections. **(C)** Representative fluorescent images of tissue sections showing neutrophil morphologies and their interactions with macrophages in granuloma subtypes. Red arrows indicate elongated neutrophils in a type I granuloma. In the ENR of type II granuloma, the yellow arrow shows the interaction between a neutrophil and an infected macrophage, the white arrow shows an infected macrophage containing phagocytosed neutrophil debris, and the white box shows a macrophage engulfing a whole neutrophil. Fluorescent images are 20 μm maximum projections. Images are representative of tissue sections from 2 animals, scale bar, 50 µm. Boundaries of the necrotic cores in fluorescent images are marked with white dashed lines.

**Fig. S4. scRNA-seq analysis of mature granulomas reveals neutrophil subgroups with distinct transcript profiles. (A)** UMAP plot of scRNA-seq data of neutrophils from micro-dissected granulomas. (**B-D)** Manhattan plots depicting gene ontology (GO) analysis of transcript profiles from the neutrophil subgroups shown in **(A).** GO terms relevant to this study are highlighted and numbered. MF – Molecular Function, BP – Biological Process, KEGG - Kyoto Encyclopedia of Genes and Genomes **(E)** Expression maps of representative genes in neutrophil subgroups, with the corresponding neutrophil subgroups marked by black dashed lines. **(B-D)** False Discovery Rate (FDR) correction was used to determine the adjusted p-values.

**Fig. S5. Characterization of Rac2D57N fish during *M. marinum* infection. (A)** Representative bright-field and fluorescent images of WT and Rac2D57N larval fish showing neutrophil (green) recruitment to the hindbrain ventricle (outlined with dotted lines) at 3 dpi with *M .marinum* (red). Scale bar, 100 µm. Fluorescent images are maximum projections spanning the entire hindbrain ventricle. **(B)** Mean number of neutrophils recruited to the hindbrain ventricle of WT and Rac2D57N larvae following mock or *M .marinum* infection (3dpi). Each data point represents the number of neutrophils recruited to the hindbrain ventricle of a single larva. WT: 41 fish; Rac2D57N: 36 fish; WT mock: 14 fish; Rac2D57N mock: 12 fish. One-way ANOVA with Tukey’s multiple comparison test was used. **(C) i & ii:** Mean number of neutrophils observed in the Rac2D57N granulomas and their WT siblings. Each data point represents the number of neutrophils observed in a single granuloma. The data shows the remaining two independent experiments corresponding to Fig. 2I. **i)** n = 41 granulomas for WT & 37 granulomas for the Rac2D57N, obtained from 3 animals each; **ii)** n = 30 granulomas for both WT and Rac2D57N, obtained from 3 animals each. **i)** Two-tailed, unpaired t-test; **ii)** Unpaired t-test with Welch’s correction. **(D)** Fluorescent images of granuloma explants showing altered neutrophil morphologies in Rac2D57N-expressing neutrophils (white arrows in the insets) compared to a WT neutrophil (red arrow). Scale bar, 50 µm. **(E) i & ii:** *M. marinum* burden in WT and Rac2D57N granulomas represented as mean arbitrary fluorescence units (AU). Granulomas were dissected from 14 dpi animals. Each data point represents *M. marinum* fluorescence measured in a single granuloma. The data shows the remaining two independent experiments corresponding to Fig. 2K. **i)** n = 40 granulomas for WT & 37 granulomas for the Rac2D57N, obtained from 3 animals each; **ii)** n = 16 granulomas for both WT and Rac2D57N, obtained from 3 animals each. **i & ii)** Unpaired t-test with Welch’s correction. **(B, C, E)** Error bars show SD.

**Fig. S6. *devR* regulon plays a key role in neutrophil-driven modulation of mycobacterial burden. (A)** Bacterial burden in WT *M. marinum* and *devR* mutant-infected granulomas dissected from WT fish, represented as mean arbitrary fluorescence units (AU). For WT *M. marinum,* n = 36 granulomas; for *devR* mutant, n = 41 granulomas. Each data point represents mycobacterial fluorescence measured in a single granuloma, with granulomas obtained from 3 animals per group. Error bars indicate SD. Unpaired t-test with Welch’s correction. The data shows the second independent experiment corresponding to Fig. 4G. **(B)** Granuloma bacterial burden in WT and Rac2D57N fish infected with either WT *M. marinum* (red data points) or *M. marinum* overexpressing *devR* & *devS* (cyan data points), represented as mean arbitrary fluorescence units (AU). For WT *M. marinum* infection: n = 39 granulomas each from WT and Rac2D57N fish. For *M. marinum devRS* overexpressor infection: n = 40 granulomas from WT fish and 41 granulomas from Rac2D57N fish. Brown-Forsythe and Welch ANOVA tests, followed by Dunnett’s T3 multiple comparisons test, were used. Granulomas were dissected from 14 dpi animals. Each data point represents mycobacterial fluorescence measured in a single granuloma, with granulomas obtained from 3 animals per group. Error bars indicate SD. The data shows the second independent experiment corresponding to Fig. 4I. **(C)** Model describing the role of neutrophils in promoting granuloma mycobacterial survival through the regulation of the *devR* regulon. Neutrophils release an unknown signal, either through diffusion or cell death in the early necrotic region. This signal is sensed by extracellular mycobacteria via the membrane-associated gas sensor, histidine protein kinase DevS, part of a two-component regulatory system. DevS then undergoes autophosphorylation, leading to the phosphorylation and activation of the response regulator DevR, a DNA-binding protein that drives the expression of the *devR* regulon. This regulon enhances mycobacterial survival under the hostile conditions present in necrotic granulomas, such as hypoxia and NO stress. The unknown signal may act cumulatively with known DevS signals, like hypoxia and NO, to upregulate the *devR* regulon. The artwork was created with BioRender.com.

**Fig. S7. Identification of host genes differentially regulated in neutrophil-deficient granulomas. (A)** Principal-component analysis (PCA) of the host transcriptome from *M. marinum-*infected wild-type and Rac2D57N granulomas. (B) Volcano plot showing differential expression of the host genes in Rac2D57N vs WT granulomas. The horizontal dotted line indicates a p_adj_ threshold of 0.1, whereas the vertical dotted lines represent log2 fold change thresholds of −1 and 1 respectively. **(C)** Bubble plot based on Gene Set Enrichment Analysis (GSEA) of the host transcriptome, showing Hallmark gene sets from the Molecular Signatures Database (MSigDB) that are up or downregulated in Rac2D57N vs WT granulomas. **(D)** GSEA bubble plot of the host transcriptome, showing the top ten Curated gene sets from MSigDB that are significantly up or downregulated in Rac2D57N vs WT granulomas. **(C, D)** The size of each circle represents the weighted number of genes involved in the term. The color and intensity of the bubbles represent the enrichment score and -log10 (p_adj_) values, respectively. Gene sets relevant to this study are marked in red. **(E, F, G)** Enrichment score (ES) plots for the gene sets corresponding to TNF-α signaling, E2F targets, and NRF2 targets, as highlighted in C & D. Positive and negative enrichment scores indicate gene sets that are over-represented among the most upregulated or downregulated genes in Rac2D57N granulomas. Black vertical bars represent individual genes within a gene set, with their position reflecting each gene’s contribution to the overall enrichment score. Genes appearing at or before the enrichment score maximum (**E, G**) and at or after the enrichment minimum (**F**) contribute to the enrichment signal. The bottom portion of the plot shows the value of the ranking metric along the list of ranked genes. **(H, I, J)** Heatmaps showing relative expression levels for the host genes associated with TNF-α signaling, E2F targets, and NRF2 targets from WT and Rac2D57N granulomas.

**Fig. S8. Identification of additional mycobacterial transcripts enriched in necrotic wild-type granulomas. (A)** Heatmaps showing mycobacterial genes specifically upregulated in zebrafish necrotic granulomas vs in vivo mouse macrophages, related to lipid metabolism, deoxyribonucleotide biosynthesis (*nrd* operon), lipoproteins, and transcriptional regulators. **(B)** Box plots showing the median log normalized counts of representative transcripts from **(A)** in WT granulomas and broth culture. Each data point represents the log normalized count of the respective transcript from an individual experiment. For the fatty acid shuttle, *M. marinum* shows upregulation of *echA6* but not the functional analog *fabH* in granulomas suggesting a key role for *echA6* in fatty acid transfer and cell envelope lipid biosynthesis in this niche. In the interrelated PDIM-PGL biosynthesis pathways, specific upregulation of the *fadD26* compared to genes involved in PGL precursor synthesis suggests a shift in the balance towards PDIM production in granulomas. Benjamini-Hochberg (BH) correction was used to determine the adjusted p-values.

## Supplementary movie legends

**Movie S1.** Split-screen video showing neutrophils (green) tracked over 230 minutes in *M. marinum* (red) infected type I and extra-necrotic region of type II granuloma explants. Frames were captured at 10-minute intervals. Individual tracks are labeled in white, magenta, and red. Maximum intensity projections, 100 μm. Scale bar, 50 µm. The video corresponds to **Fig. 1E**.

**Movie S2.** Neutrophil migration between sub-compartments of a type II granuloma. The movement of two neutrophils (red and white arrows), initially found near the necrotic core, toward the *M. marinum* containing extra-necrotic region (white box) was tracked over 240 mins. Frames were captured at 10-minute intervals. Maximum intensity projection, 100 μm. Scale bar, 50 µm. The video corresponds to **Fig. S1**.

**Movie S3.** Neutrophil viability in *M. marinum* infected type I and extra-necrotic region (white box) of type II granulomas. Granulomas were observed for 790 minutes, with individual frames captured at 10-minute intervals. Maximum intensity projections, 100 μm. Scale bar, 50 µm. The video corresponds to **Fig. 1G**.

**Movie S4.** Video showing the viability of an uninfected neutrophil (red arrow) and an infected neutrophil (white arrow) in the *M. marinum* containing extra-necrotic region of a type II granuloma. Individual frames were acquired at 10-minute intervals. 100 μm maximum intensity projection. Scale bar, 50 µm. The video corresponds to **Fig. S2**.

**Movie S5.** Split-screen video showing neutrophil (green) and macrophage (red) viability in *M. marinum* (magenta) infected type I and extra-necrotic region (white box) of type II granulomas. Granulomas were observed for 780 minutes, with frames captured at 10-minute intervals. 100 μm maximum intensity projections. Scale bar, 50 µm. The video corresponds to **Fig. S3A**.

**Movie S6.** Interaction between a dying neutrophil (red arrow) and an *M. marinum* (magenta) infected macrophage (white arrow) in the extra-necrotic region of a type II granuloma. Neutrophils are labeled green and macrophages in red. The yellow arrow indicates *M. marinum* released from the infected macrophage following its interaction with the neutrophil. Frames were captured at 10-minute intervals. Maximum intensity projection, 100 μm. Scale bar, 25 µm. The video corresponds to **Fig. S3B**.

**Movie S7.** Neutrophils (green) tracked over 230 minutes in *M. marinum* (red) infected granuloma explants, 5 hours post-treatment with DMSO/duvelisib. Frames were captured at 10-minute intervals. Individual tracks are labeled in white, magenta, and red. 100 μm maximum intensity projections. Scale bar, 50 µm. The video corresponds to **Fig. 2C**.

## Supplementary table legends

**Table S1.** ScRNA-seq data showing the proportion of granuloma neutrophil subgroups and differentially expressed genes within each subgroup

**Table S2.** Gene Ontology (GO) terms enriched in each neutrophil subgroup

**Table S3.** Sequences of zebrafish and *M. marinum* rRNA depletion probes used in the dual RNA-seq experiment in this study

**Table S4.** RNA-seq data showing differentially expressed *M. marinum* genes in Rac2D57N vs WT granulomas

**Table S5.** RNA-seq data showing differentially expressed zebrafish genes in Rac2D57N vs WT granulomas. Separate tabs present Gene Set Enrichment Analysis (GSEA) data for the zebrafish transcriptome, including Hallmark and Curated (C2) gene sets from the Molecular Signatures Database (MSigDB) that are upregulated or downregulated in Rac2D57N vs WT granulomas. Additionally, expression data for the host genes related to key gene sets, such as TNF-α signaling, E2F targets, and NRF2 targets are provided separately.

**Table S6.** RNA-seq data showing differentially expressed *M. marinum* genes in WT granulomas vs broth culture.

**Table S7.** Expression data for mycobacterial genes commonly upregulated in zebrafish necrotic granulomas and in vivo mouse macrophages.

**Table S8.** Expression data for mycobacterial genes specifically upregulated in zebrafish necrotic granulomas compared to in vivo mouse macrophages.

**Table S9.** Lineage-specific non-synonymous and synonymous mutations identified in the top 50 granuloma-specific genes. Genes with multiple non-synonymous mutations are highlighted in green. The Sequence Read Archive (SRA) list of 69 clinical isolates is provided in a separate tab.

