## Supplementary material for "Granuloma Dual RNA-Seq Reveals Composite Transcriptional Programs Driven by Neutrophils and Necrosis within Tuberculous Granulomas": Combined Supplementary Figures

**Fig. S1**

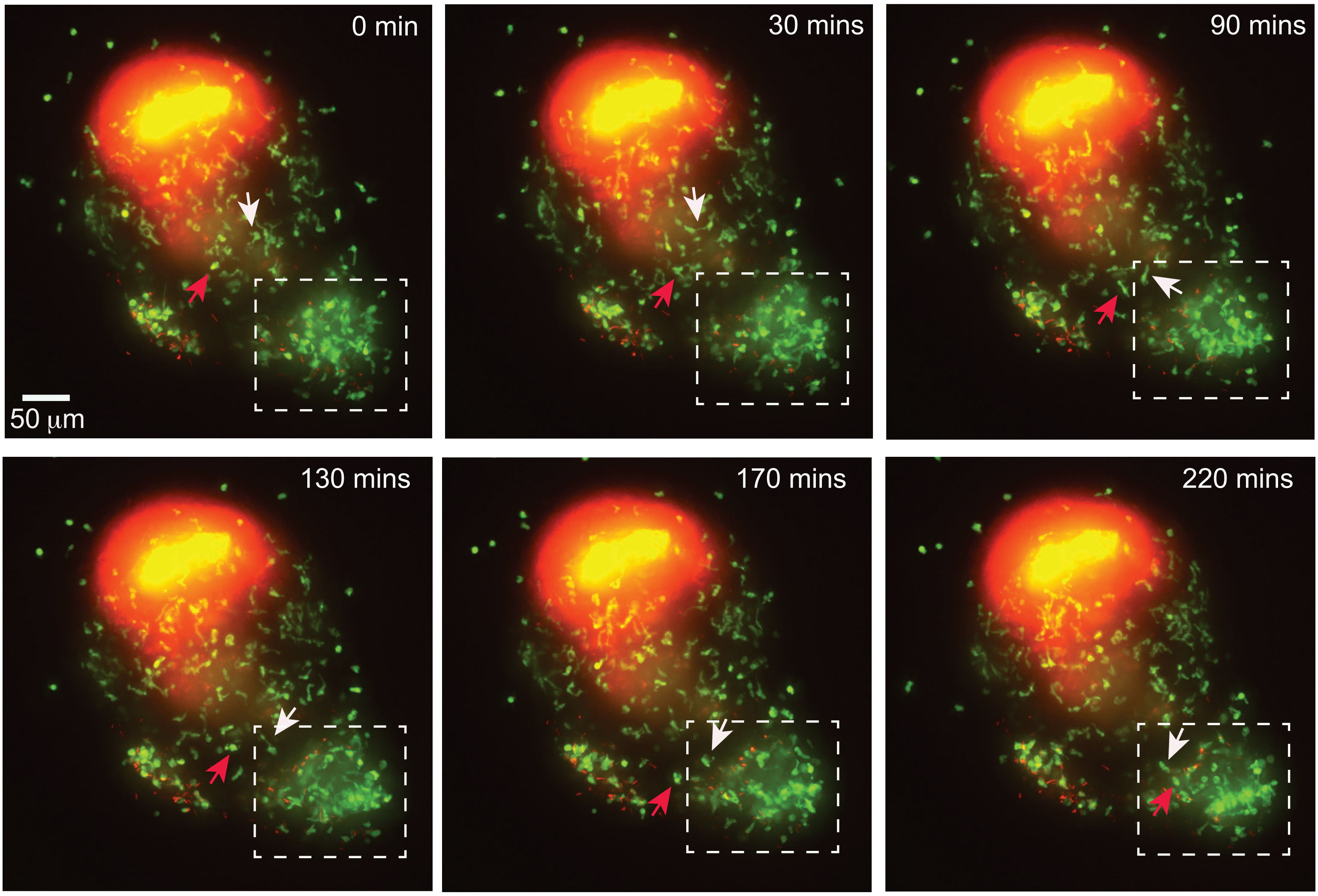

**Fig. S2**

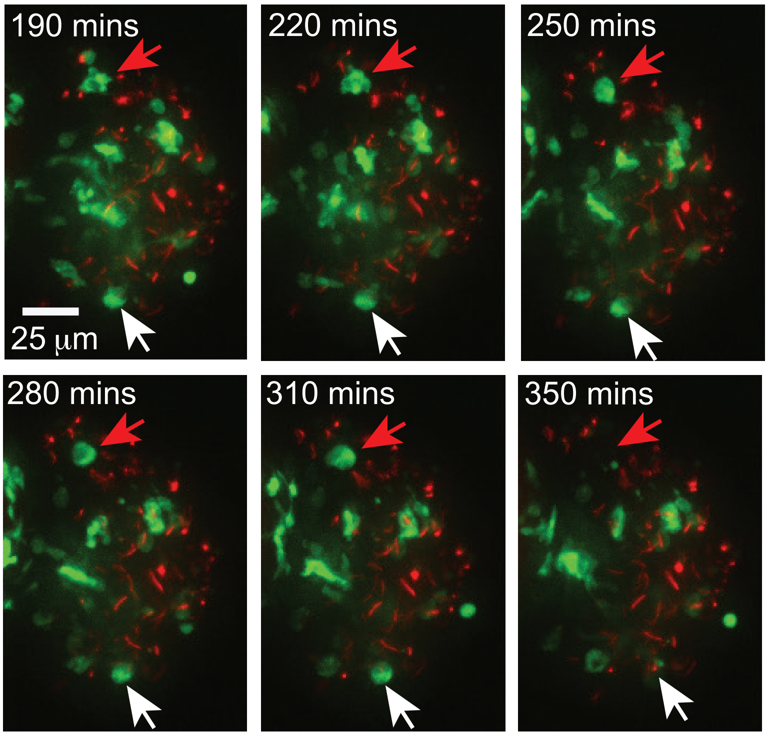

Fig. S3

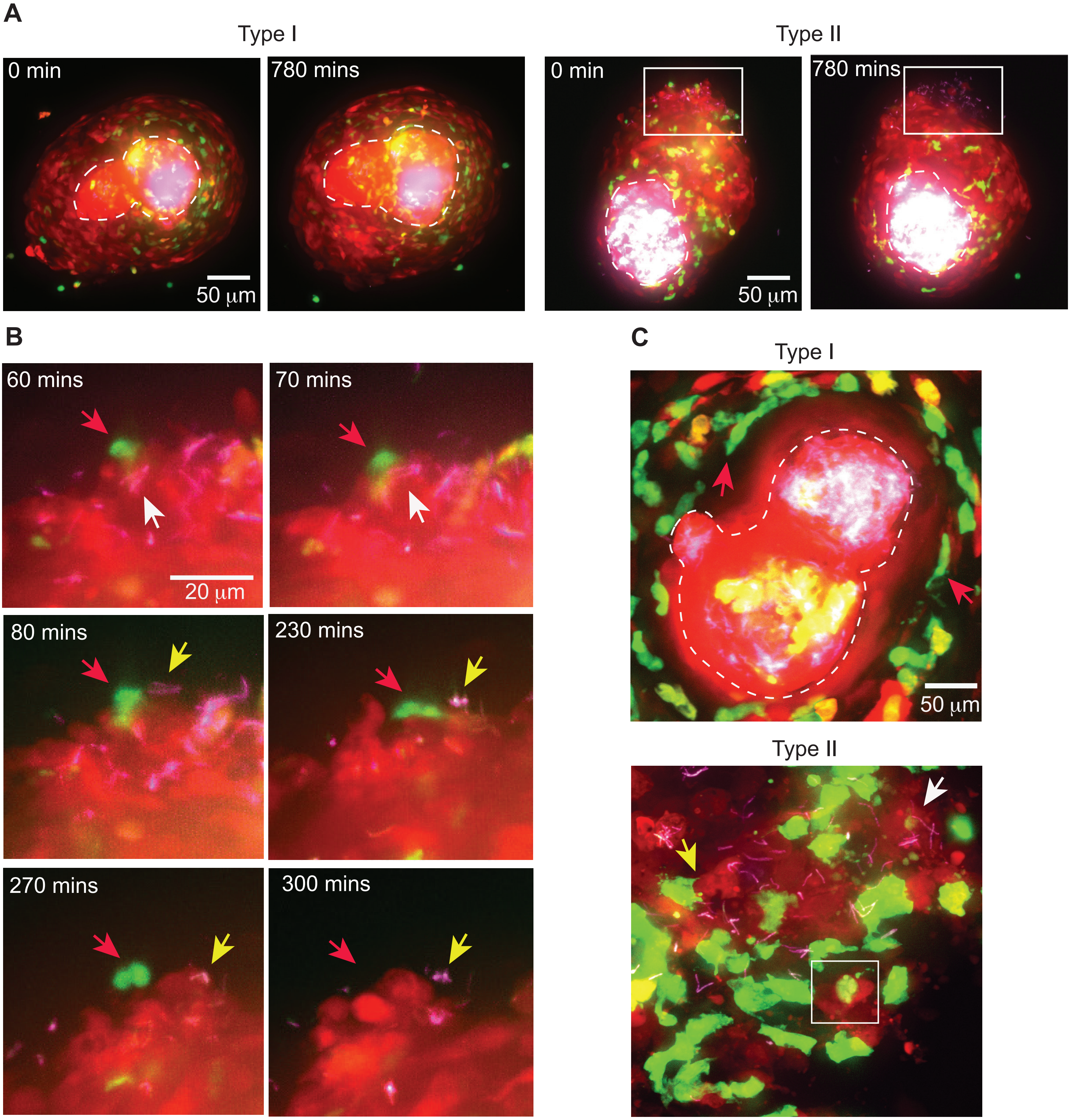
A
Type I
0 min
780 mins
50 μm
Type II
0 min
780 mins
50 μm
B
60 mins
70 mins
20 μm
80 mins
230 mins
270 mins
300 mins
C
Type I
50 μm
Type II

Fig. S4

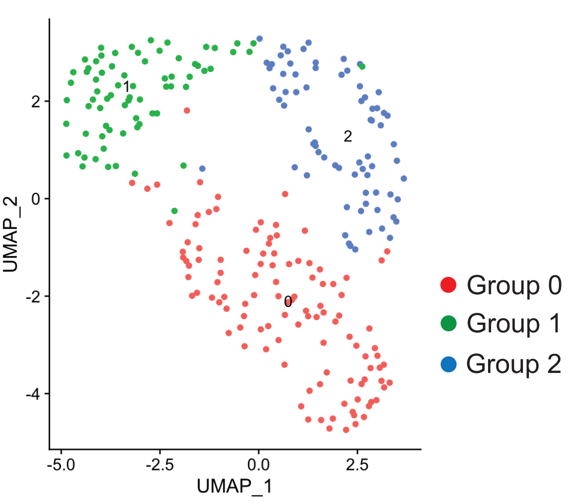


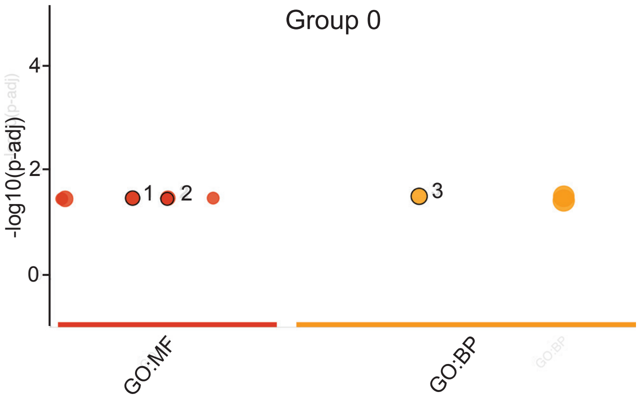


| id | source | term_id | term_name | term_size | | p_value |
|---|---|---|---|---|---|---|
| 1 | GO:MF | GO:0023026 | MHC class II protein complex binding | 7 | | 3.5e-02 |
| 2 | GO:MF | GO:0042287 | MHC protein binding | 5 | | 3.6e-02 |
| 3 | GO:BP | GO:0019882 | antigen processing and presentation | 12 | | 3.2e-02 |

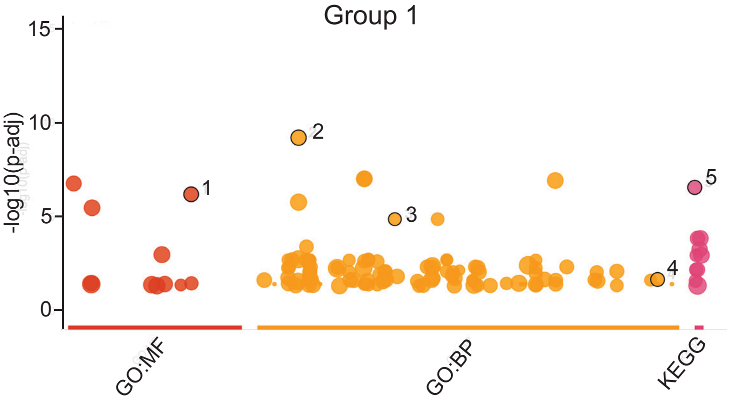


| id | source | term_id | term_name | term_size | | p_value |
|---|---|---|---|---|---|---|
| 1 | GO:MF | GO:0051015 | actin filament binding | 93 | | 6.7e-07 |
| 2 | GO:BP | GO:0007015 | actin filament organization | 152 | | 6.7e-10 |
| 3 | GO:BP | GO:0034314 | Arp2/3 complex-mediated actin nucleation | 23 | | 1.4e-05 |
| 4 | GO:BP | GO:1990266 | neutrophil migration | 46 | | 2.2e-02 |
| 5 | KEGG | KEGG:00010 | Glycolysis / Gluconeogenesis | 56 | | 2.9e-07 |

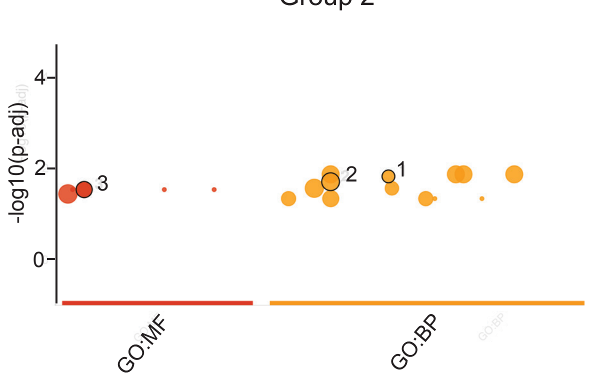


| id | source | term_id | term_name | term_size | | p_value |
|---|---|---|---|---|---|---|
| 1 | GO:BP | GO:0030223 | neutrophil differentiation | 14 | | 1.5e-02 |
| 2 | GO:BP | GO:0009605 | response to external stimulus | 638 | | 2.0e-02 |
| 3 | GO:MF | GO:0005125 | cytokine activity | 149 | | 3.0e-02 |

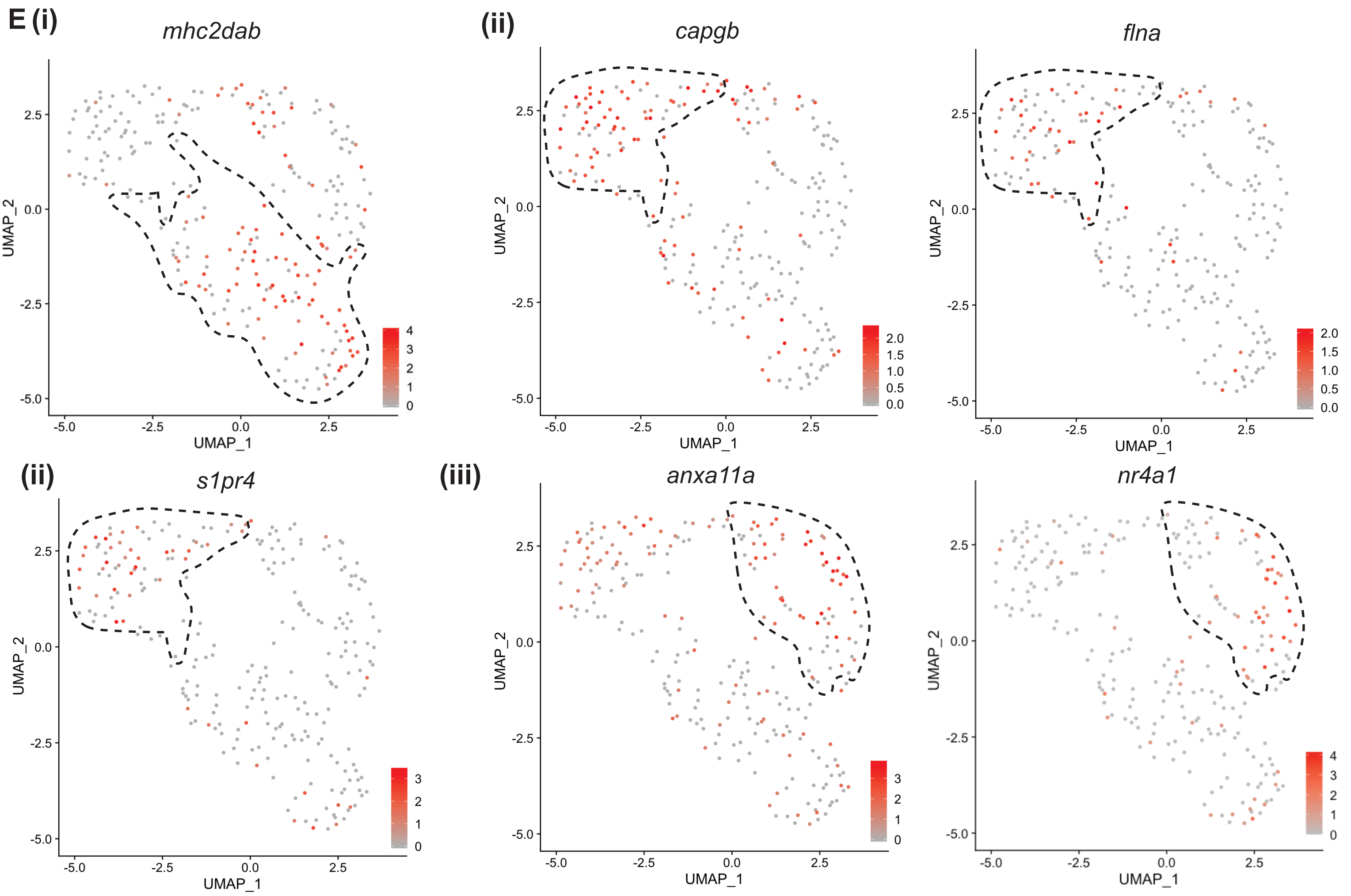

Fig. S5

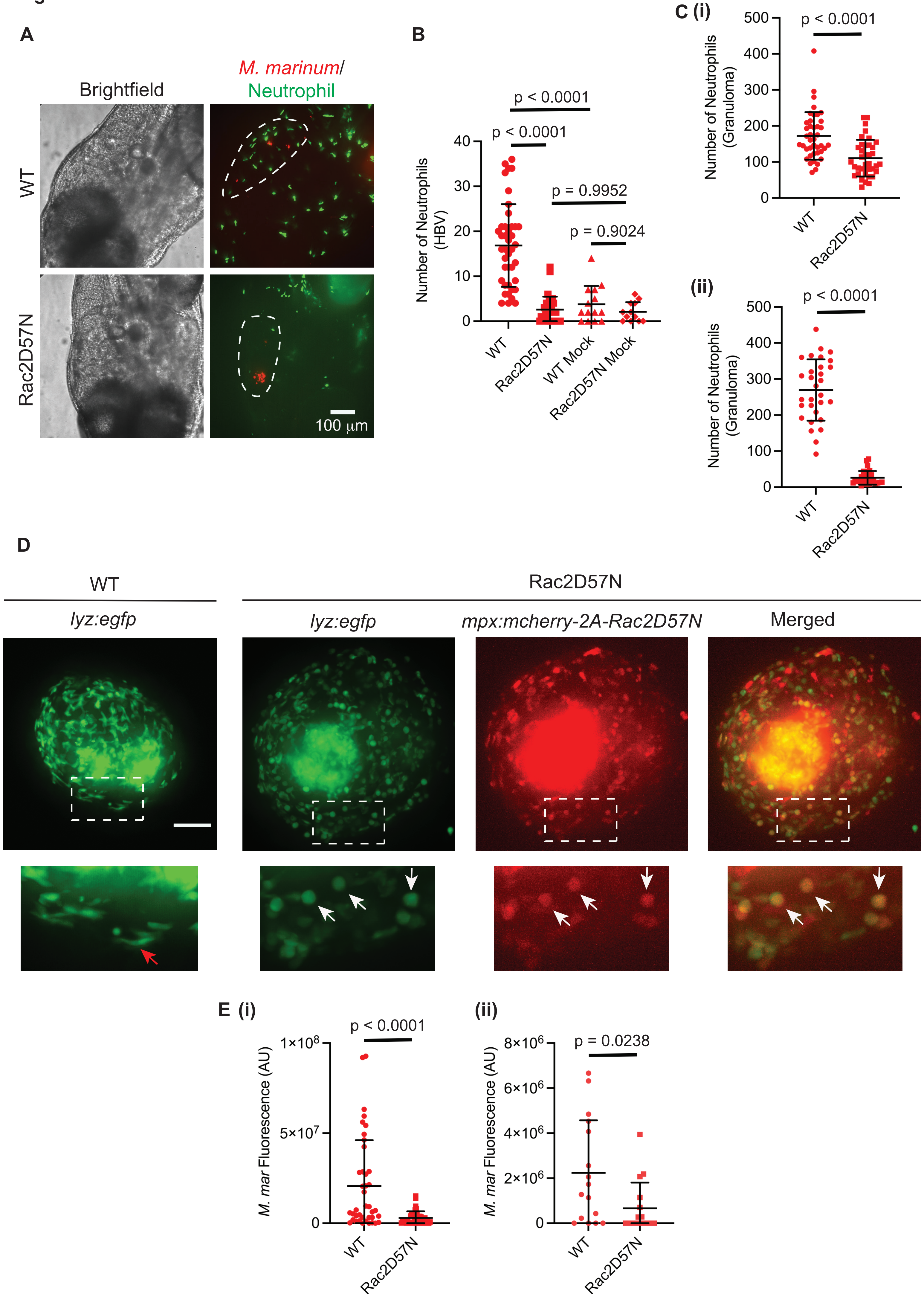

A
Brightfield
M. marinum/
Neutrophil
WT
Rac2D57N
100 μm
B
p < 0.0001
p < 0.0001
p = 0.9952
p = 0.9024
Number of Neutrophils (HBV)
WT
Rac2D57N
WT Mock
Rac2D57N Mock
C (i)
p < 0.0001
Number of Neutrophils (Granuloma)
WT
Rac2D57N
(ii)
p < 0.0001
Number of Neutrophils (Granuloma)
WT
Rac2D57N
D
WT
Rac2D57N
lyz:egfp
lyz:egfp
mpx:mcherry-2A-Rac2D57N
Merged
E (i)
p < 0.0001
M. mar Fluorescence (AU)
1×10^8
5×10^7
WT
Rac2D57N
(ii)
p = 0.0238
M. mar Fluorescence (AU)
8×10^6
6×10^6
4×10^6
2×10^6
WT
Rac2D57N

**Fig. S6**

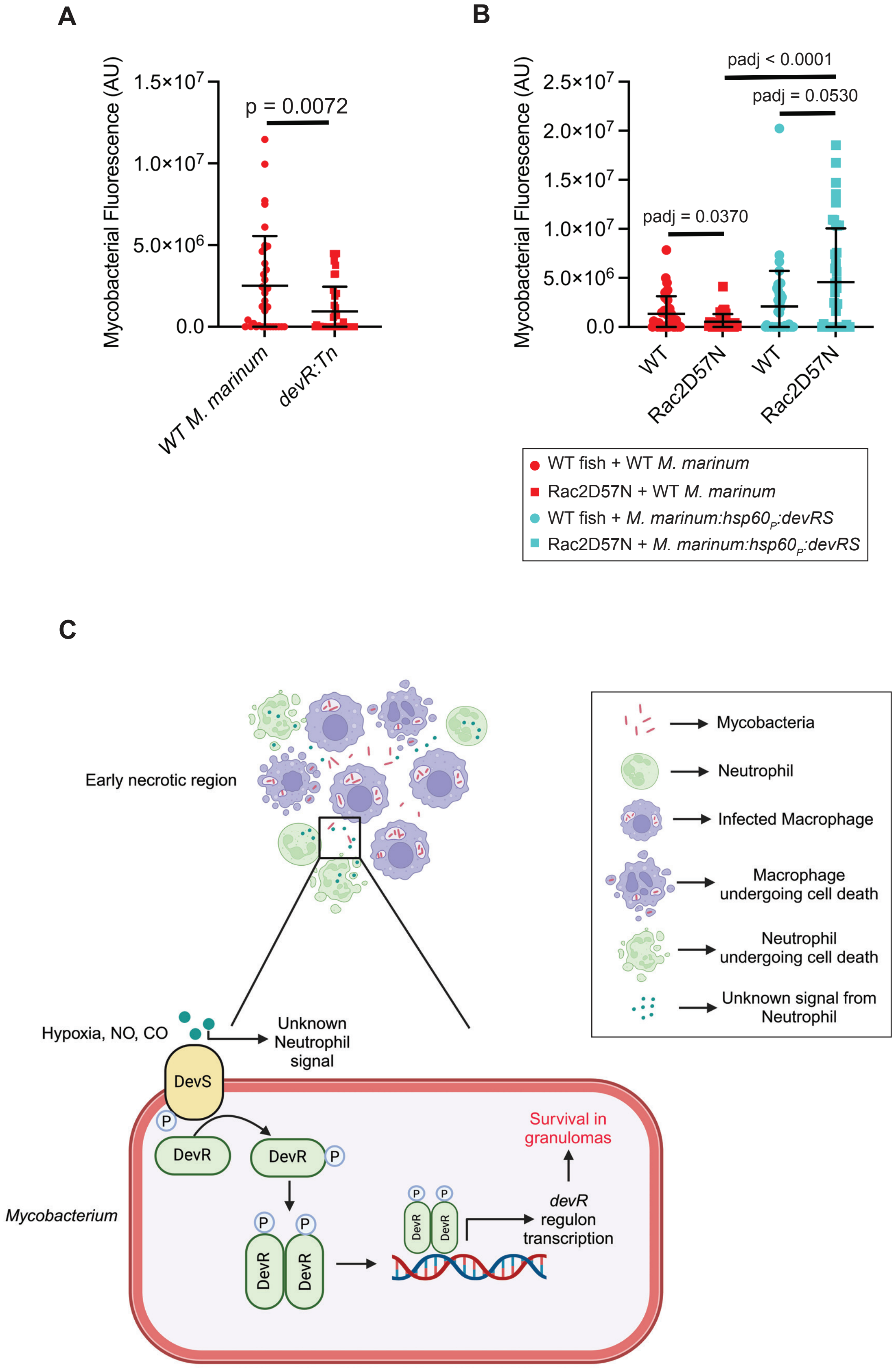
A
Mycobacterial Fluorescence (AU)
1.5×10^7
1.0×10^7
5.0×10^6
0.0
p = 0.0072
WT M. marinum
devR:Tn
B
Mycobacterial Fluorescence (AU)
2.5×10^7
2.0×10^7
1.5×10^7
1.0×10^7
5.0×10^6
0.0
padj < 0.0001
padj = 0.0530
padj = 0.0370
WT
Rac2D57N
WT
Rac2D57N
WT fish + WT M. marinum
Rac2D57N + WT M. marinum
WT fish + M. marinum:hsp60P:devRS
Rac2D57N + M. marinum:hsp60P:devRS
C
Early necrotic region
Mycobacteria
Neutrophil
Infected Macrophage
Macrophage undergoing cell death
Neutrophil undergoing cell death
Unknown signal from Neutrophil
Hypoxia, NO, CO
Unknown Neutrophil signal
DevS
P
DevR
DevR
P
P
P
DevR
DevR
P
P
DevR
DevR
devR regulon transcription
Survival in granulomas
Mycobacterium

Fig. S7

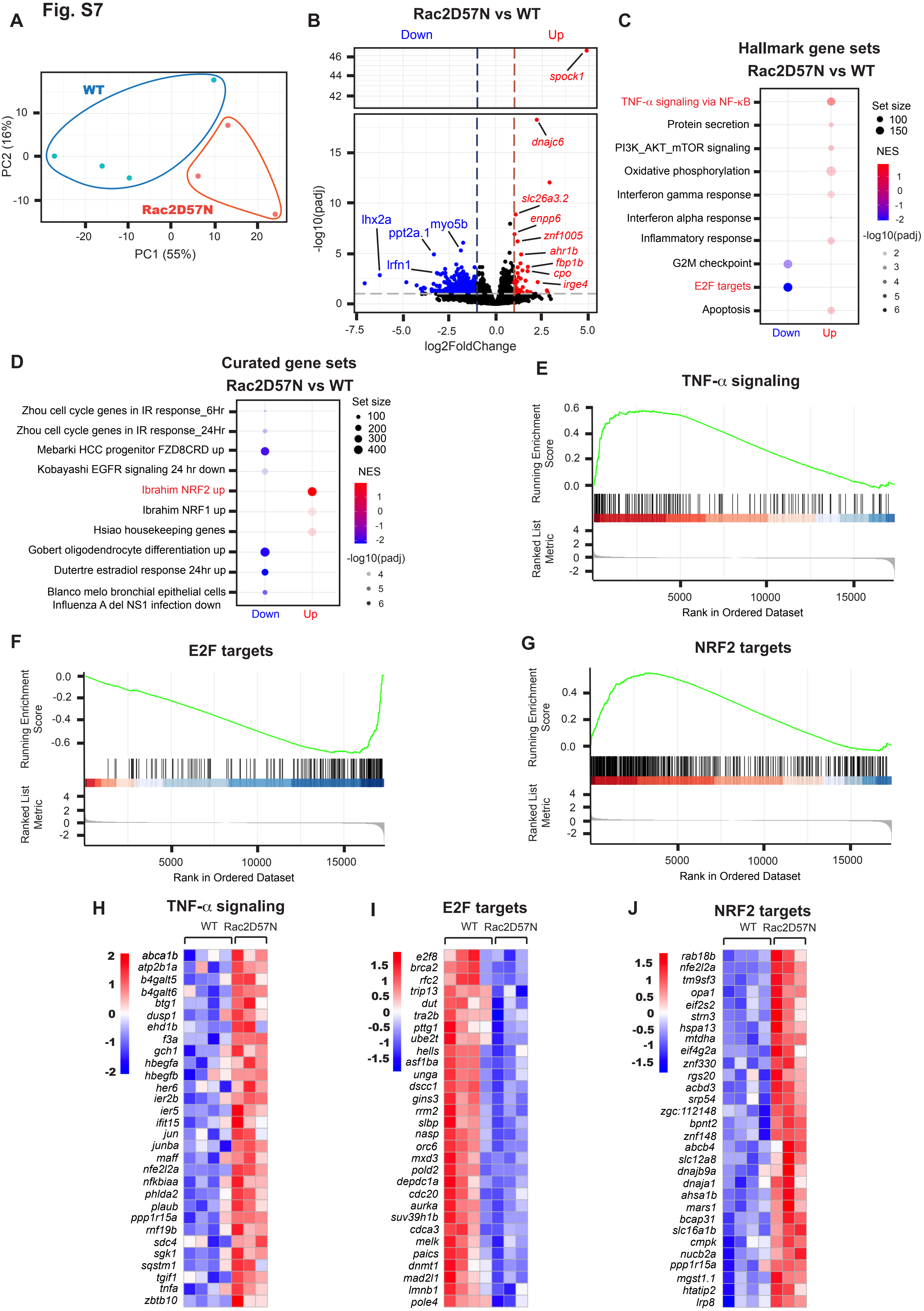
A
WT
Rac2D57N
PC2 (16%)
PC1 (55%)
B
Rac2D57N vs WT
Down
Up
spock1
dnajc6
slc26a3.2
enpp6
znf1005
ahr1b
fbp1b
cpo
irge4
lhx2a
ppt2a.1
myo5b
lrfn1
-log10(padj)
log2FoldChange
C
Hallmark gene sets
Rac2D57N vs WT
TNF-α signaling via NF-κB
Protein secretion
PI3K_AKT_mTOR signaling
Oxidative phosphorylation
Interferon gamma response
Interferon alpha response
Inflammatory response
G2M checkpoint
E2F targets
Apoptosis
Down
Up
Set size
NES
-log10(padj)
D
Curated gene sets
Rac2D57N vs WT
Zhou cell cycle genes in IR response_6Hr
Zhou cell cycle genes in IR response_24Hr
Mebarki HCC progenitor FZD8CRD up
Kobayashi EGFR signaling 24 hr down
Ibrahim NRF2 up
Ibrahim NRF1 up
Hsiao housekeeping genes
Gobert oligodendrocyte differentiation up
Dutertre estradiol response 24hr up
Blanco melo bronchial epithelial cells
Influenza A del NS1 infection down
Down
Up
Set size
NES
-log10(padj)
E
TNF-α signaling
Running Enrichment Score
Ranked List Metric
Rank in Ordered Dataset
F
E2F targets
Running Enrichment Score
Ranked List Metric
Rank in Ordered Dataset
G
NRF2 targets
Running Enrichment Score
Ranked List Metric
Rank in Ordered Dataset
H
TNF-α signaling
WT
Rac2D57N
abca1b
atp2b1a
b4galt5
b4galt6
btg1
dusp1
ehd1b
f3a
gch1
hbegfa
hbegfb
her6
ier2b
ier5
ifit15
jun
junba
maff
nfe2l2a
nfkbiaa
phlda2
plaub
ppp1r15a
rnf19b
sdc4
sgk1
sqstm1
tgif1
tnfa
zbtb10
I
E2F targets
WT
Rac2D57N
e2f8
brca2
rfc2
trip13
dut
tra2b
pttg1
ube2t
hells
asf1ba
unga
dscc1
gins3
rrm2
slbp
nasp
orc6
mxd3
pold2
depdc1a
cdc20
aurka
suv39h1b
cdca3
melk
paics
dnmt1
mad2l1
lmnb1
pole4
J
NRF2 targets
WT
Rac2D57N
rab18b
nfe2l2a
tm9sf3
opa1
eif2s2
strn3
hspa13
mtdha
eif4g2a
znf330
rgs20
acbd3
srp54
zgc:112148
bpnt2
znf148
abcb4
slc12a8
dnajb9a
dnaja1
ahsa1b
mars1
bcap31
slc16a1b
cmpk
nucb2a
ppp1r15a
mgst1.1
htatip2
lrp8

Fig. S8

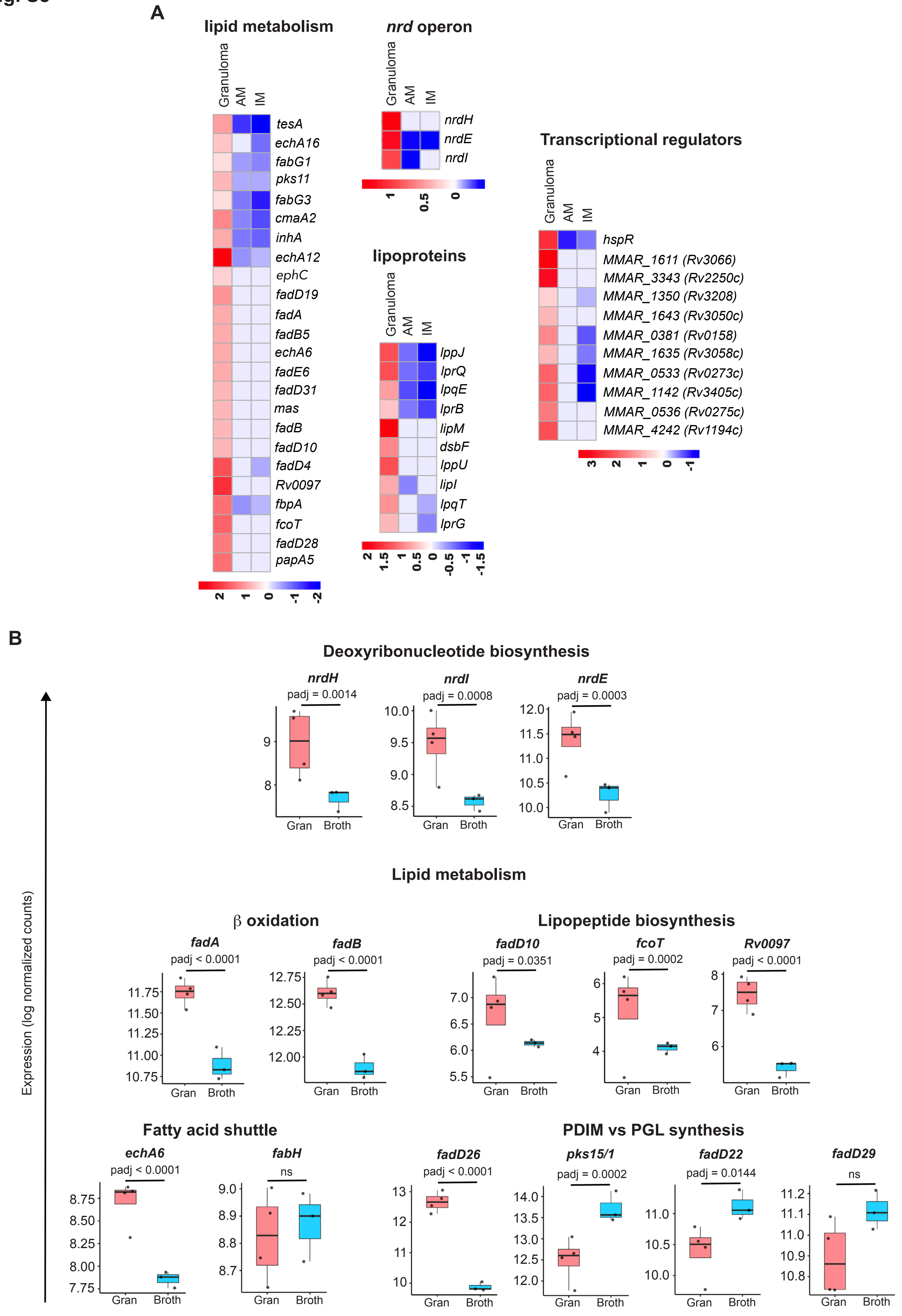
A
lipid metabolism
Granuloma
AM
IM
tesA
echA16
fabG1
pks11
fabG3
cmaA2
inhA
echA12
ephC
fadD19
fadA
fadB5
echA6
fadE6
fadD31
mas
fadB
fadD10
fadD4
Rv0097
fbpA
fcoT
fadD28
papA5
2
1
0
-1
-2
nrd operon
nrdH
nrdE
nrdI
1
0.5
0
lipoproteins
lppJ
lprQ
lpqE
lprB
lipM
dsbF
lppU
lipI
lpqT
lprG
2
1.5
1
0.5
0
-0.5
-1
-1.5
Transcriptional regulators
hspR
MMAR_1611 (Rv3066)
MMAR_3343 (Rv2250c)
MMAR_1350 (Rv3208)
MMAR_1643 (Rv3050c)
MMAR_0381 (Rv0158)
MMAR_1635 (Rv3058c)
MMAR_0533 (Rv0273c)
MMAR_1142 (Rv3405c)
MMAR_0536 (Rv0275c)
MMAR_4242 (Rv1194c)
3
2
1
0
-1
B
Expression (log normalized counts)
Deoxyribonucleotide biosynthesis
nrdH
padj = 0.0014
nrdI
padj = 0.0008
nrdE
padj = 0.0003
Gran
Broth
Lipid metabolism
β oxidation
fadA
padj < 0.0001
fadB
padj < 0.0001
Lipopeptide biosynthesis
fadD10
padj = 0.0351
fcoT
padj = 0.0002
Rv0097
padj < 0.0001
Fatty acid shuttle
echA6
padj < 0.0001
fabH
ns
PDIM vs PGL synthesis
fadD26
padj < 0.0001
pks15/1
padj = 0.0002
fadD22
padj = 0.0144
fadD29
ns
